# Heterotypic responses against nsp12/nsp13 from prior SARS-CoV-2 infection associates with lower subsequent endemic coronavirus incidence

**DOI:** 10.1101/2023.10.23.563621

**Authors:** David J. Bean, Janet Monroe, Yan Mei Liang, Ella Borberg, Yasmeen Senussi, Zoe Swank, Sujata Chalise, David Walt, Janice Weinberg, Manish Sagar

**Affiliations:** Department of Virology, Immunology and Microbiology, Boston University Chobanian & Avedisian School of Medicine, Boston, MA; Department of Medicine, Boston University Chobanian & Avedisian School of Medicine, Boston, MA; Department of Pathology, Brigham and Women’s Hospital, Harvard Medical School, Boston, MA; Wyss Institute for Biologically Inspired Engineering, Harvard University, Boston, MA; Department of Biostatistics, Boston University School of Public Health, Boston, MA

## Abstract

Immune responses from prior SARS-CoV-2 infection and COVID-19 vaccination do not prevent re-infections and may not protect against future novel coronaviruses (CoVs). We examined the incidence of and immune differences against human endemic CoVs (eCoV) as a proxy for response against future emerging CoVs. Assessment was among those with known SARS-CoV-2 infection, COVID-19 vaccination but no documented SARS-CoV-2 infection, or neither exposure. Retrospective cohort analyses suggest that prior SARS-CoV-2 infection, but not COVID-19 vaccination alone, protects against subsequent symptomatic eCoV infection. CD8^+^ T cell responses to the non-structural eCoV proteins, nsp12 and nsp13, were significantly higher in individuals with previous SARS-CoV-2 infection as compared to the other groups. The three groups had similar cellular responses against the eCoV spike and nucleocapsid, and those with prior spike exposure had lower eCoV-directed neutralizing antibodies. Incorporation of non-structural viral antigens in a future pan-CoV vaccine may improve protection against future heterologous CoV infections.

## Introduction

There are seven known human coronaviruses (HCoV) with consequences ranging from asymptomatic infection or mild respiratory symptoms to severe respiratory distress or death.^1^ Four (HCoV-229E, HCoV-NL63, HCoV-OC43, and HCoV-HKU1) of these, termed endemic CoVs (eCoVs), mostly cause only mild symptomatic illness and are responsible for 15-30% of the “common cold” in human adult.^2,3^ However, in the past twenty years, three HCoVs have emerged that are highly pathogenic and significantly more deadly: Severe Acute Respiratory Syndrome (SARS-CoV); Middle East Respiratory Syndrome (MERS); and SARS-CoV-2, responsible for the coronavirus disease 2019 (COVID-19) pandemic.^4,5^ Betacoronavirus, one of four coronavirus (CoV) genera, includes several of the most clinically relevant HCoVs, including HCoV-OC43, HCoV-HKU1, SARS-CoV, SARS-CoV-2, and MERS-CoV.^6^

Numerous SARS-CoV-2 vaccines, primarily targeting the spike protein, are highly effective in reducing the incidence of hospitalization and progression to severe disease after infection.^7–9^ The SARS-CoV-2 vaccines or prior SARS-CoV-2 infection, however, are less likely to prevent breakthrough or re-infection especially with the most recent lineages, such as the Omicron variants and subvariants.^10,11^ Numerous studies have demonstrated that antibodies generated either from previous vaccination, prior SARS-CoV-2 infection, or both have lower neutralization potency against the Omicron variants and the subvariants.^12–14^ The less potent neutralizing antibodies (nAbs) potentially account for the limited protection against subsequent symptomatic infections among those with prior SARS-CoV-2 exposure via vaccination or infection.

Besides nAbs, cellular responses are also likely important in reducing the incidence of symptomatic infection and onset of severe disease.^15–17^ NAbs exclusively target the SARS-CoV-2 spike, while T cells respond to various peptides present across the entire SARS-CoV-2 coding genome.^18,19^ In general, spike sequences are more variable than other parts of the genome, both when comparing different SARS-CoV-2 variants and the diverse CoVs.^20,21^ Furthermore, in contrast to nAbs, adaptive T cell responses show slower decay over time.^22,23^ Thus, cellular responses generated via vaccination and or infection may be especially important in protecting against the development of symptomatic disease by the SARS-CoV-2 variants and the other circulating CoVs.

Previous studies from multiple groups have demonstrated the pre-existence of SARS-CoV-2 nAbs and cellular responses among individuals with no known SARS-CoV-2 exposure.^24,25^ Studies from our group and others suggests that prior infection with eCoVs and the subsequent immune response can attenuate COVID-19 disease severity after SARS-CoV-2 infection.^26,27^ These studies imply that an eCoV infection may generate heterotypic immunity against other CoV viral family members. Deciphering the immune basis for this potential heterologous immunity is important because of the ongoing threat of a novel coronavirus as the etiologic agent for a future pandemic.

In this study, we examined heterotypic immunity generated from SARS-CoV-2 infection as compared to COVID-19 vaccination. We find that prior SARS-CoV-2 infection, but not COVID-19 vaccination alone, associates with protection against subsequent symptomatic disease from the eCoVs. This heterologous immunity is likely mediated via CD8^+^ T cell responses targeting non-structural proteins, such as nsp12 and nsp13, and not nAbs. Our observations have important implications for future pan-CoV vaccines and other disease prevention strategies.

## Results

### Individuals with prior SARS-CoV-2 infection have lower incidence of subsequent symptomatic documented eCoVs

We examined the incidence of symptomatic eCoV and non-CoV infections documented on comprehensive respiratory panel PCR (CRP-PCR) tests among individuals who presented for clinical evaluation during the study period of November 30, 2020 to October 8, 2021 at Boston Medical Center (BMC) (Figure 1A). The CRP-PCR test detects the four eCoVs, SARS-CoV-2, and sixteen other common non-CoV respiratory pathogens. All individuals were categorized into three groups based on their pre-CRP-PCR test history: 1) prior documented SARS-CoV-2 infection; 2) antecedent COVID-19 vaccination but no known SARS-CoV-2 infection; 3) no previous SARS-CoV-2 antigen exposure. There were 5,713 CRP-PCR results from 4,935 (median 1 test per person, range 1 - 8) different individuals. As expected, the individuals with no prior SAR-CoV-2 exposure were younger and healthier compared to the people in the other two groups because documented SARS-CoV-2 infection occurred in and COVID-19 vaccines were preferentially given to older individuals with pre-existing comorbidities (Supplementary Table 1). In general, the differences in age, gender, and the proportion with pre-existing diagnoses was smaller among those with previous documented SARS-CoV-2 infection and those with prior COVID-19 vaccination but no known SARS-CoV-2 infection (Supplementary Table 1).

**Figure 1.**
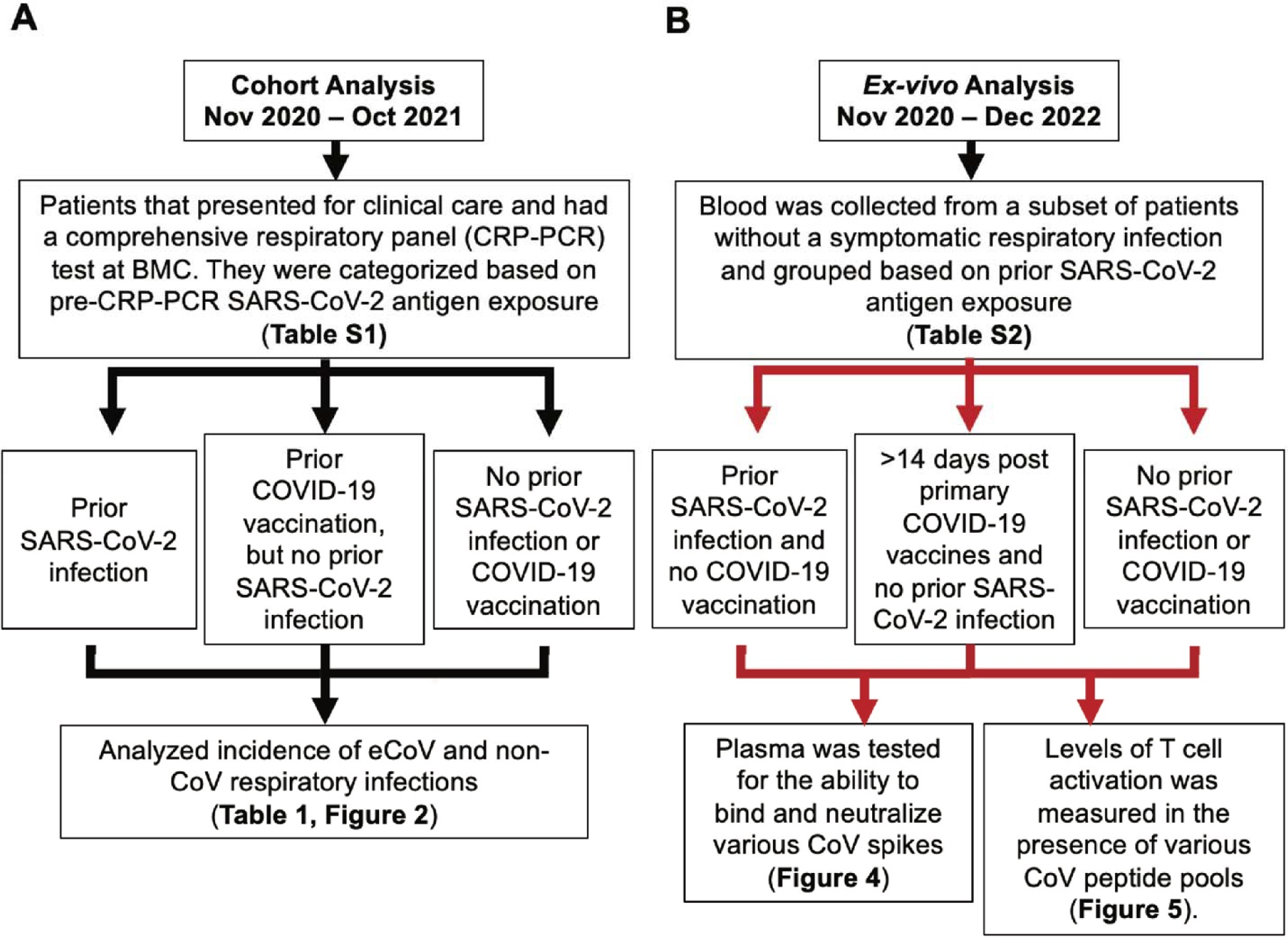
Schematic of the overall study design and workflow. (A) The chart (black arrows) details how cohort was grouped and analyzed for the incidence of respiratory infections at Boston Medical Center (BMC). (B) The chart (red arrows) details the recruitment and grouping of individuals for the ex-vivo assessment of CoV-specific immunity.

Among the 4,935 individuals, 617 had a non-SARS-CoV-2 respiratory pathogen detected on the CRP-PCR test at some time during the study period (Table 1). There were 103 eCoVs, and rhinovirus/enterovirus (n = 409) was the most common non-CoV infection detected on the CRP-PCR tests (Table 1). Of the eCoV infections, 79 were HCoV-OC43, 19 were HCoV-229E, and 5 were HCoV-NL63. Individuals with prior SARS-CoV-2 infection had the lowest incidence of subsequent symptomatic eCoV as compared to two other groups as judged by outcome alone (Table 1). Those with prior documented SARS-CoV-2 infection had lower incidence of both alpha and beta eCoVs, although only the difference in alpha eCoVs was statistically significant (Table 1). On the other hand, non-CoV and enterovirus/rhinovirus detection was not different in the three groups (Table 1). Time to event analysis also showed that eCoV (p < 0.0001, log-rank test, Figure 2A) but not non-CoV (p = 0.2823, log-rank test, Figure 2B) infections occurred less frequently among those with prior documented SARS-CoV-2 infections as compared to the individuals with COVID-19 vaccination or no known antigen exposure.

**Table 1.** Infections detected on CRP-PCR tests.

|  | Prior SARS-CoV-2 infection (n = 501) | Prior COVID-19 vaccine / No SARS-CoV-2 infection (n = 1,565) | No prior SARS-CoV-2 exposure (n = 2,869) | p-value<sup>A</sup> |
| --- | --- | --- | --- | --- |
| **Abnormal CRP-PCR<sup>B</sup>** | 51 (10.2) | 226 (14.4) | 340 (11.9) | 0.0113 |
| **eCoV** | 5 (1.0) | 45 (2.9) | 53 (1.8) | 0.0145 |
| **alpha-eCoV<sup>C</sup>** | 0 (0.0) | 13 (0.8) | 11 (0.4) | 0.0316 |
| **beta-eCoV<sup>D</sup>** | 5 (1.0) | 32 (2.0) | 42 (1.5) | 0.1777 |
| **eCoV – full vaccine<sup>E</sup>** | 3/209 (1.4) | 44/1463 (3.0) | N/A | 0.2638<sup>G</sup> |
| **eCoV – partial vaccine<sup>F</sup>** | 0/17 (0.0) | 1/102 (1.0) | N/A | 0.9999<sup>G</sup> |
| **eCoV – no vaccine** | 2/275 (0.7) | N/A | 53/2869 (1.8) | 0.2301<sup>G</sup> |
| **Non-CoV** | 46 (9.2) | 181 (11.6) | 287 (10.0) | 0.1688 |
| **Rhinovirus/Enterovirus** | 31 (6.2) | 127 (8.1) | 251 (8.7) | 0.1518 |

<sup>A</sup> Chi-Square test unless noted otherwise
<sup>B</sup> One of the twenty non-SARS-CoV-2 respiratory pathogens
<sup>C</sup> Includes HCoV-229E and HCoV-NL63
<sup>D</sup> Includes HCoV-OC43 and HCoV-HKU-1
<sup>E</sup> Received at least two doses of the Pfizer BioNTech BNT162b2 or Moderna mRNA-1273 COVID-19 vaccine or one dose of the Janssen / Johnson & Johnson Ad26.COVS2 COVID-19 vaccine a minimum of 14 days prior to the CRP-PCR test.
<sup>F</sup> Received a vaccine dose but did not meet definition of fully vaccinated as above
<sup>G</sup> Fisher's exact test
N/A: not applicable

**Figure 2.**
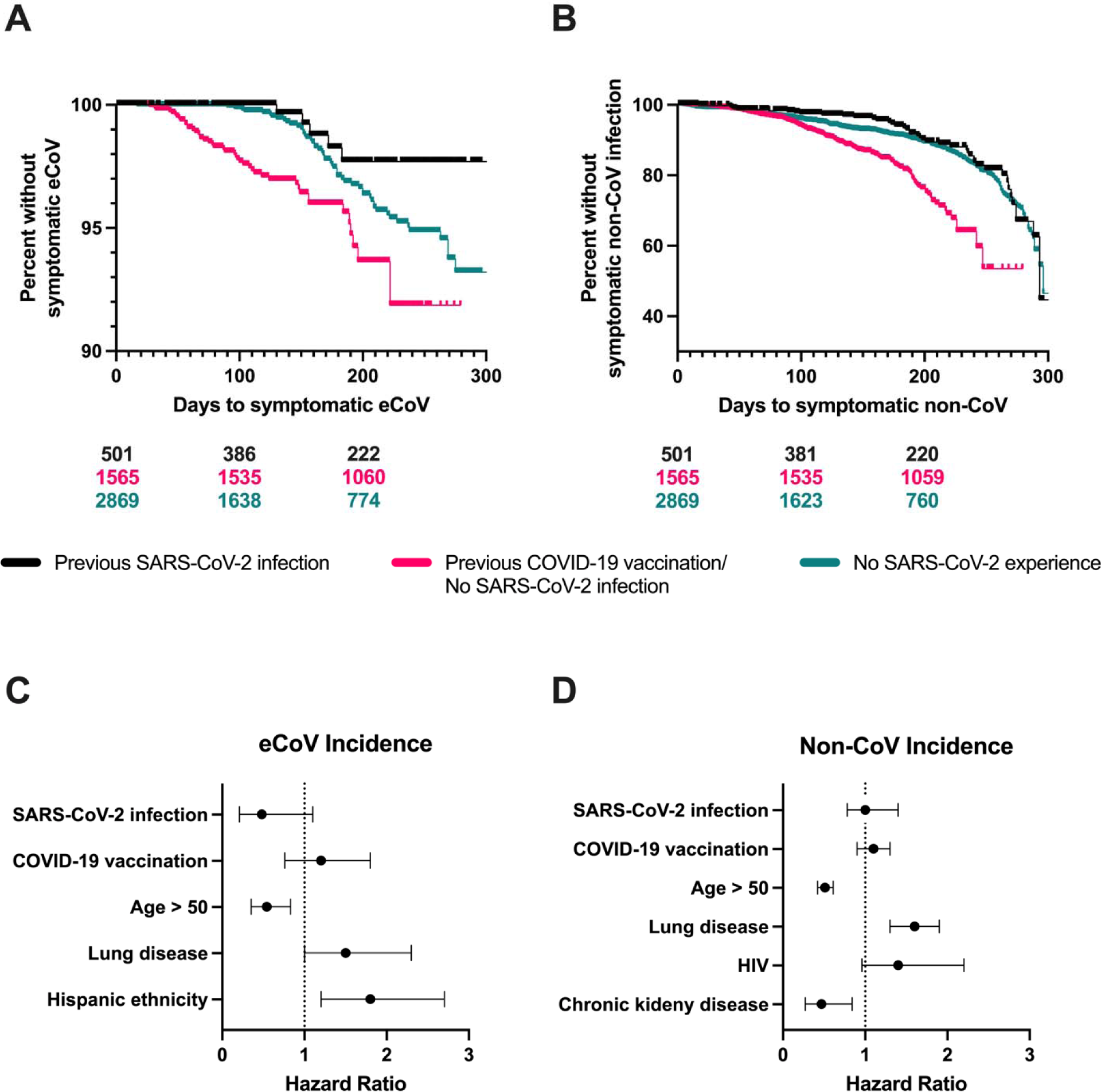
Incidence of eCoV and non-CoV infections among those with different SARS-CoV-2 antigen exposure. Kaplan Meier plot for eCoV (A) and non-CoV (B) in those with no prior documented SARS-CoV-2 antigen exposure (teal), prior COVID-19 vaccination but no known SARS-CoV-2 infection (pink), and prior SARS-CoV-2 infection (black). Plots show unadjusted analyses with number of individuals at risk in each group at the bottom. Hazard ratio (HR) of eCoV (C) and non-CoV (D) incidence in time varying adjusted models. The figures show the covariates that had a p-value ≤ 0.1 in the multivariable model. No prior documented SARS-CoV-2 antigen exposure group is the reference category. The horizontal dotted lines indicate the HR for each variable along with the 95% confidence interval, and the vertical dotted line indicates a HR of 1.0.

Among those with prior COVID-19 vaccination but no documented SARS-CoV-2 infection, 1,463 of the 1,565 (93%) were deemed fully vaccinated because they had received the last dose of the recommended primary vaccine series at least fourteen days before the CRP-PCR result. Among the 501 individuals with prior SARS-CoV-2 infection, 226 (45%) had at least one COVID-19 vaccine dose antecedent to the subsequent CRP-PCR test (Table 1). Importantly, a symptomatic eCoV infection was significantly lower in those with prior SARS-CoV-2 infection and no vaccination (2 of 275, 0.7%) as compared to the individuals that had been deemed fully vaccinated but had no known prior SARS-CoV-2 infection (44 of 1,463, 3.0%, p = 0.0245, Fischer’s exact test, Table 1).

We also conducted time-varying covariate analysis because some individuals had SARS-CoV-2 infection or COVID-19 vaccination during and not prior to the study period. In these instances, the CRP-PCR test of interest also occurred after the exposure. Importantly, this time-varying multivariable model accounted for baseline demographics, such as age, gender, and pre-existing comorbidities (Supplementary Table 1). Previous COVID-19 infection (hazard ratio (HR) 0.48, 95% confidence interval (CI) 0.21, 1.1, p=0.08) associated with around 50% reduced risk of a future symptomatic eCoV (Figure 2C), but it did not impact non-CoV incidence (Figure 2D). On the other hand, COVID-19 vaccination showed no effect on the risk for eCoV or non-CoV infections. There was no statistically significant interaction between prior SARS-CoV-2 infection and antecedent COVID-19 vaccination in these models. Interestingly, age greater than 50 years had significantly lower eCoV (HR 0.54, 95% CI 0.35, 0.83, p = 0.01) and non-CoV (HR 0.51, 95% CI 0.42, 0.61, p < 0.0001) incidence in these models. The lower hazard for both infections suggests decreased overall exposure to respiratory viruses in this older age group. Those with preexisting lung disease, however, had higher risk for both eCoVs (HR 1.0, 95% CI 1.0, 2.3, p = 0.04) and non-CoVs (HR 1.6, 95% CI 1.3, 1.9, p < 0.0001) infection as potentially expected.

### Ex-vivo examination of immunological differences among the groups

We collected blood samples between November 2020 to December 2022 to examine immune differences associated with the observed protection against symptomatic eCoVs. In contrast to the retrospective analysis, individuals in this independent cohort were grouped into three categories with stricter definitions (Figure 1B). First, none of those with prior PCR documented SARS-CoV-2 infection had COVID-19 vaccination (n= 20). Second, all the individuals in the COVID-19 vaccine group had completed the recommended primary vaccine series at least 14 days prior to blood collection (n = 25). Finally, individuals with no known SARS-CoV-2 antigen exposure (n = 28) were interviewed regarding prior suspected infection and COVID-19 vaccination during the consent process before blood collection. We specifically did not choose pre-pandemic samples for the no antigen exposure group because eCoV circulation was significantly higher in the preceding years as compared to the year when COVID-19 was declared a public health emergency in the city of Boston.^28^ Thus, samples collected prior to as compared to after pandemic onset may have higher eCoV immune responses from greater community virus circulation. The three groups had relatively similar demographics other than age distribution (Table S2). Additionally, the duration between the last documented exposure to SARS-CoV-2 spike was shorter among the individuals in the COVID-19 vaccination as compared to the SARS-CoV-2 infection group (Table S2).

The individuals classified as not having a previous SARS-CoV-2 infection may have had prior undiagnosed or asymptomatic COVID-19.^29^ Thus, there is a possibility of misclassification. We used a comprehensive antibody and T-cell based assessments to evaluate this possibility. First, we measured anti-SARS-CoV-2 (receptor binding domain (RBD) and nucleocapsid) plasma IgG levels by the Simoa SARS-CoV-2 IgG antibody test.^30^ RBD antibody response has been demonstrated as highly indicative of SARS-CoV-2 spike exposure either through infection or vaccination.^31^ As expected, individuals with prior SARS-CoV-2 spike exposure via infection or vaccination had higher RBD antibody levels compared to those with no known antigen exposure (Figure 3A). Nucleocapsid IgG reactivity was used to further classify some COVID-19 vaccine recipients that may have had prior SARS-CoV-2 infection.^32,33^ Again as expected, nucleocapsid IgG levels were significantly higher in individuals with documented SARS-CoV-2 infection as compared to the two other groups (Figure 3B). We also examined the number of activated T cells (both CD4^+^ and CD8^+^) after exposure to various SARS-CoV-2 peptide pools using the activation induced marker (AIM) assay (Supplementary Figure 1). As expected, higher percentage of activated CD4^+^ (CD134^+^ CD137^+^, Figure 3C) and CD8^+^ (CD69^+^ CD137^+^, Figure 3D) T cells were observed after SARS-CoV-2 spike peptide pool stimulation in individuals with either a previous SARS-CoV-2 infection or COVID-19 vaccination compared to those individuals without a known history of SARS-CoV-2 exposure. The activated SARS-CoV-2 spike specific CD4^+^ and CD8^+^ T cell levels were similar between individuals with either a previous SARS-CoV-2 infection or COVID-19 vaccination. Also as expected, individuals with a previous documented SARS-CoV-2 infection had elevated SARS-CoV-2 nucleocapsid responsive CD4^+^ (Figure 3E) and CD8^+^ (Figure 3F) T cells compared to individuals with COVID-19 vaccination only or no history of SARS-CoV-2 antigen experience.

**Figure 3.**
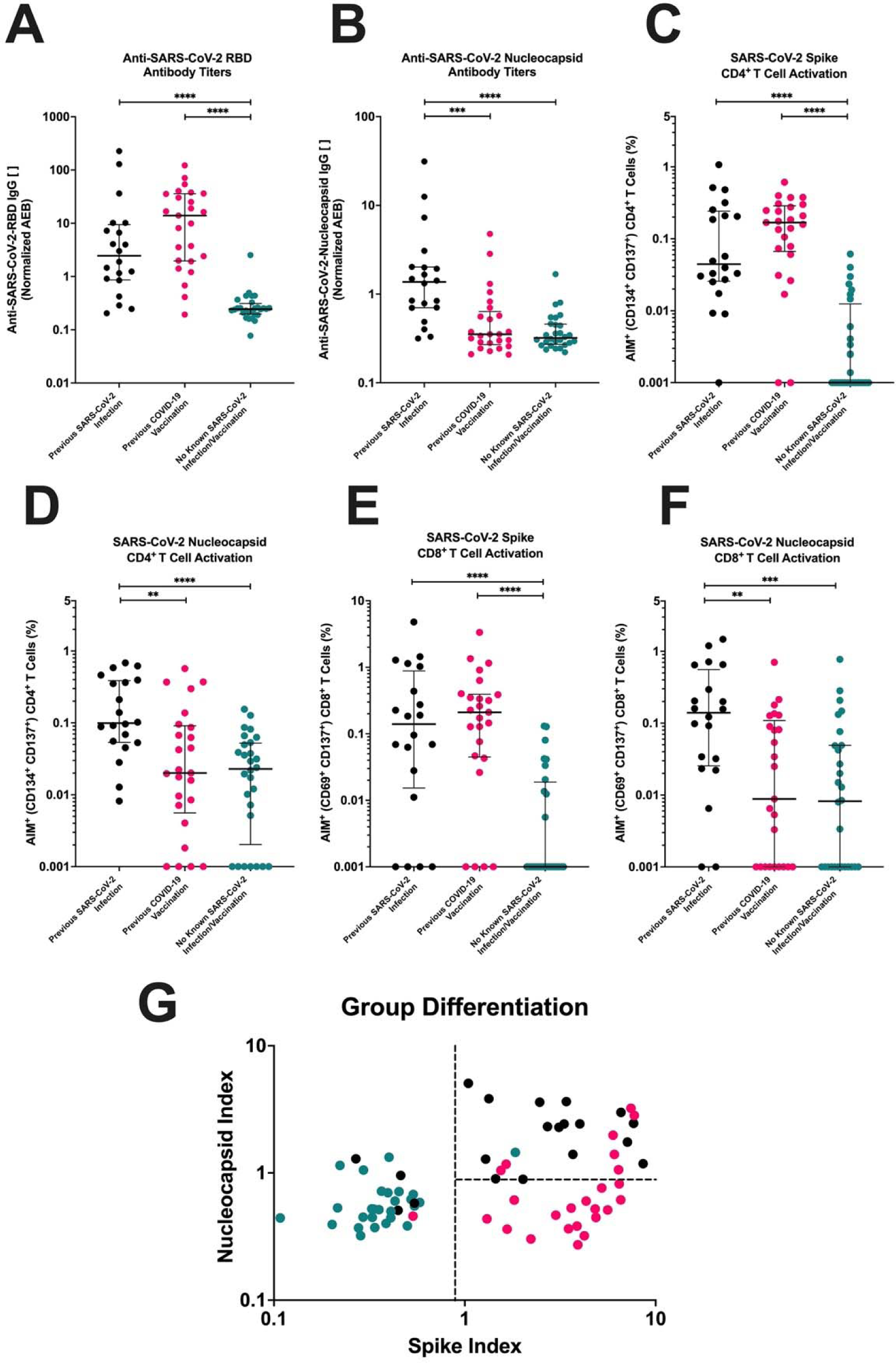
Differentiation of the type of SARS-CoV-2 antigen exposure based on adaptive immune responses to SARS-CoV-2 antigens. Adaptive immune responses against SARS-CoV-2 spike (A,C,E) and SARS-CoV-2 nucleocapsid (B,D,F) in those with no prior SARS-CoV-2 exposure (teal), prior COVID-19 vaccination but no SARS-CoV-2 infection (pink), and prior documented SARS-CoV-2 infection (black). (A-B) IgG antibody levels that bound SARS-CoV-2 RBD (A) or SARS-CoV-2 nucleocapsid (B) coated beads were measured by the enzyme based Simoa detection system. Antibody concentrations were calculated based on the average enzyme per bead (AEB). (C-F) AIM assay with SARS-CoV-2 spike (C, E) or SARS-CoV-2 nucleocapsid (D, F) peptide pools and activation was measured on CD4^+^ (CD134^+^ CD137^+^) and CD8^+^ (CD69^+^ CD137^+^) T cells by flow cytometry. Data were background subtracted against the negative control (DMSO only). The dark horizontal lines in each scatter dot plot denote the median and interquartile range. Note, the y-axis varies among the different panels. (G) The antibody and T cell responses against SARS-CoV-2 (A-F) were log transformed and combined to create a spike and nucleocapsid index. The dotted lines represent the cutoff values to best differentiate the groups. **, ***, **** represent p-values <0.01, <0.001, <0.0001 respectively.

Log-transformed values from the three nucleocapsid (IgG, AIM CD4^+^ and AIM CD8^+^) and three spike (RBD, AIM CD4^+^ and AIM CD8^+^) measurements were combined to generate a nucleocapsid and spike index. Spike and nucleocapsid cutoffs were chosen that best separated the three groups (Fig. 3G). These cutoffs yielded relatively similar test characteristics and overall accuracy (82%, Table 2) compared to another method (accuracy range 84 – 90%) that was based on examining SARS-CoV-2 cellular responses only.^34^ One and seven in the no antigen exposure and COVID-19 vaccine only group respectively clustered with the majority of individuals with prior documented SARS-CoV-2 infection based on the established cutoffs. Thus, these eight individuals may have had prior undocumented or asymptomatic SARS-CoV-2 infection. In the subsequent analyses, results were also presented excluding some or all of these potentially “misclassified” individuals. Furthermore, four individuals with documented SARS-CoV-2 positive PCR results had either low spike or nucleocapsid index. Two of these four had severe COVID-19 with admission to the intensive care unit (ICU) while the other two were not hospitalized during their primary infection. Seven and nine of the remaining sixteen individuals with documented SARS-CoV-2 infection were not hospitalized and not admitted to a non-ICU floor during their disease course respectively.

**Table 2.** Accuracy of clinical classifications based on SARS-CoV-2 adaptive immune responses.

|  |  |  | Clinical Classification |  |  | Positive predictive value | Negative predicative value |
| --- | --- | --- | --- | --- | --- | --- | --- |
|  |  |  | Documented SARS-CoV-2 infection (n = 20) | COVID-19 vaccination, but no SARS-CoV-2 infection (n = 25) | No known SARS-CoV-2 antigen exposure (n = 28) |  |  |
| Cutoff Classification | Prior infection | Spike ≥ 0.89 & NC ≥ 0.89 | 16 | 7 | 1 | 67% | 92% |
|  | Vaccination, no infection | Spike > 0.89 & NC < 0.89 | 0 | 17 | 0 | 100% | 86% |
|  | No SARS-CoV-2 exposure | Spike < 0.89 | 4 | 1 | 27 | 84% | 98% |
| Sensitivity |  |  | 80% | 68% | 96% | Overall Accuracy 82% |  |
| Specificity |  |  | 85% | 100% | 89% |  |  |

### Humoral immune responses are unlikely to account for the decreased symptomatic eCoV incidence

We first investigated humoral immune responses as a factor that associated with decreased incidence of symptomatic eCoVs because nAbs have been described as the correlate of protection for SARS-CoV-2.^35,36^ We measured the ability of patient plasma to neutralize pseudoviruses that express the spike protein of various CoVs (SARS-CoV-2-Wuhan, HCoV-OC43, and HCoV-229E) (Figure 4). Even though spike exposure was more recent in those with vaccination (Table S2), neutralization of the SARS-CoV-2 spike-based pseudovirus was similar in the individuals with a previous SARS-CoV-2 infection as compared to COVID-19 vaccination even when seven COVID-19 vaccine individuals with possible misclassification were excluded (Figure 4A). As expected, individuals with SARS-CoV-2 spike exposure either via infection or vaccination had higher SARS-CoV-2 nAbs as compared to the no SARS-CoV-2 antigen exposure group (p = 0.0438, p = 0.0565 when excluding the one “misclassified” person from the no antigen group, Figure 4D).

**Figure 4.**
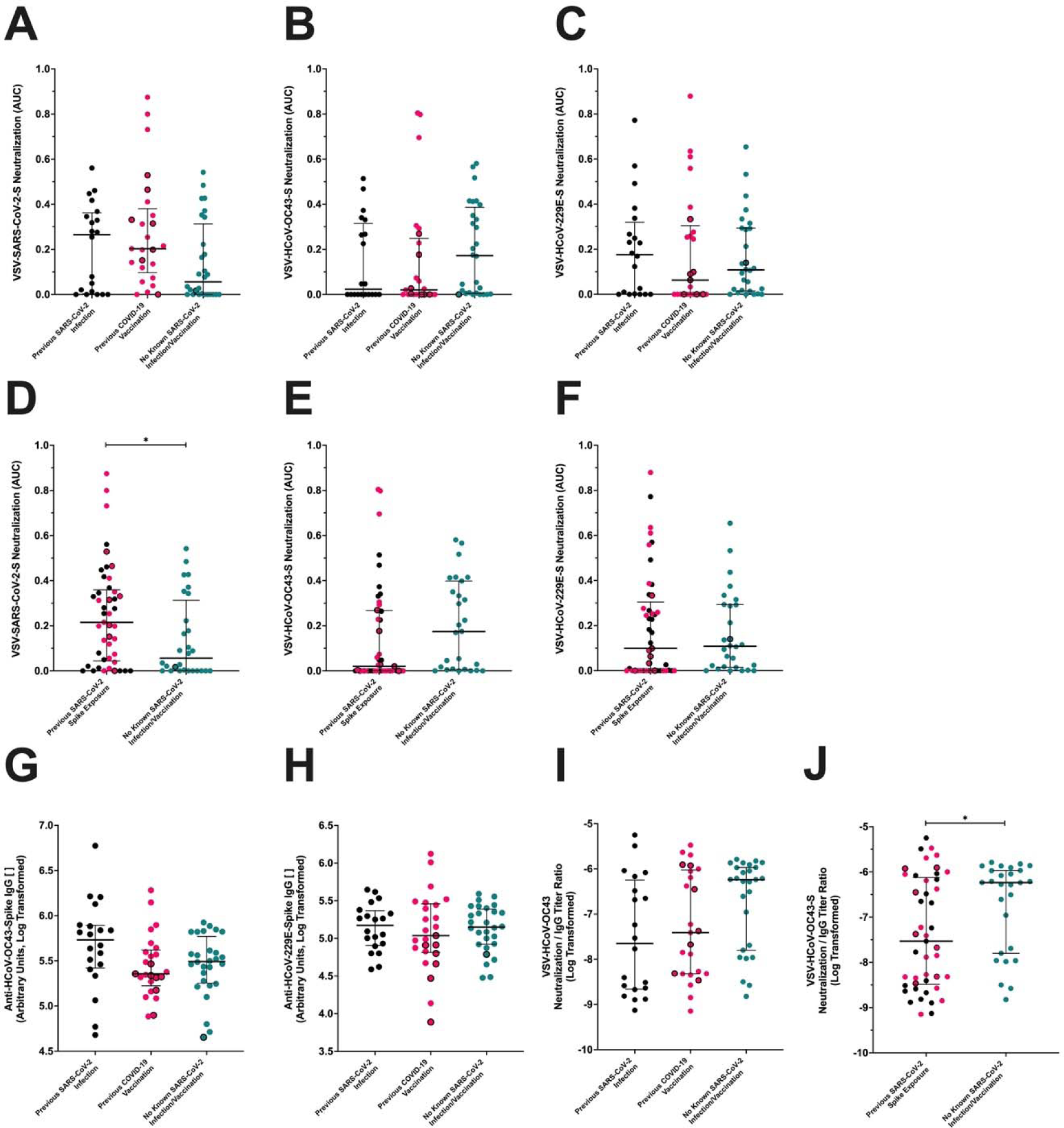
Neutralization and binding antibodies against CoV-spikes among those with different SARS-CoV-2 antigen exposure. Plasma antibody responses against various CoV spike antigens were tested in those with no prior SARS-CoV-2 exposure (teal), prior COVID-19 vaccination but no SARS-CoV-2 infection (pink), and prior documented SARS-CoV-2 infection but no COVID-19 vaccination (black). Black borders represent the eight individuals identified as potentially having prior undocumented or asymptomatic SARS-CoV-2 infection. (A-F) Area under the curve neutralization values against pseudoviruses expressing the spike proteins of SARS-CoV-2 (A, D), HCoV-OC43 (B, E), and HCoV-229E (C, F). (D-F) The same respective data as in panels A-C, but the individuals with previous SARS-CoV-2 infection and those with COVID-19 vaccination were placed in the same column because both had prior spike exposure. (G-H) ELISA derived plasma IgG levels against HCoV-OC43 (G) and HCoV-229E (H) spike. (I-J) A ratio of HCoV-OC43 specific neutralization (B) and antibody titers (G) among the three groups (I) and those with and without prior SARS-CoV-2 spike exposure (J). The dark horizontal lines in each scatter dot plot denote the median and interquartile range. * represents p-values <0.05.

No significant differences in HCoV-OC43 neutralization were observed between the prior SARS-CoV-2 infection group compared to the COVID-19 vaccinated group with or without excluding the seven people with possible prior occult infection (Figure 4B). Surprisingly, individuals with any SARS-CoV-2 spike exposure either via SARS-CoV-2 infection or COVID-19 vaccination had trending lower HCoV-OC43 neutralization responses as compared to the no SARS-CoV-2 antigen experience individuals (p = 0.0645, p = 0.0407 when excluding the one “misclassified” person from the no antigen group, Figure 4E). On the other hand, there were no differences in the ability to neutralize HCoV-229E spike-containing pseudoviruses between any of the three groups (Figure 4C) or between those with as compared to without previous spike exposure (Figure 4F).

A previous investigation had demonstrated that SARS-CoV-2 infection enhances antibodies against the beta-, but not alpha-eCoVs.^37^ We also observed that binding antibodies as measured by enzyme-linked immunosorbent assay (ELISA) were trending higher in those with documented SARS-CoV-2 infection against HCoV-OC43 (Figure 4G) but not HCoV-229E (Figure 4H). HCoV-OC43 binding antibodies were higher among those with prior SARS-CoV-2 infection in multi-variate linear regression (Table S3) and in univariate analysis when compared to the COVID-19 vaccinated group (p = 0.0449, p = 0.1494 after excluding the seven with possible prior asymptomatic infection) and the no antigen exposure group (p = 0.0702, p = 0.1001 after excluding the one “misclassified” person). Individuals with no known prior SARS-CoV-2 spike experience had the highest ratio of HCoV-OC43 neutralization relative to total binding antibodies (Fig. 4I). Importantly, the ratio of neutralization relative to total binding antibodies trended lower in those with any previous SARS-CoV-2 spike experience as compared to those with no spike antigen exposure (p = 0.0585, p = 0.0366 after excluding the one person with possible prior asymptomatic infection, Fig. 4J). Both HCoV-OC43 nAbs and ratio of HCoV-OC43 nAbs to binding antibodies, however, were not significantly different in multi-variate linear regression analysis that accounted for age, gender, and comorbidities. This suggests that SARS-CoV-2 infection and possibly COVID-19 vaccination enhances non-neutralizing binding antibodies more than nAbs against HCoV-OC43.

### HCoV-OC43 non-structural protein directed cellular immune responses are higher among those with prior SARS-CoV-2 infection

We next examined differences in cellular immune responses among the different groups using the AIM assay. We focused our testing to HCoV-OC43 peptides because of PBMC quantity limitations, and HCoV-OC43 was the primary eCoV circulating in Boston during the study period (Table 1). We also focused our study on spike, nucleocapsid, and nsp12/nsp13 T cell activity, which allowed us to compare spike, non-spike, structural, and non-structural protein responses.

Similar HCoV-OC43-reactive CD4^+^ and CD8^+^ T cells were observed between the three groups when PBMCs were stimulated with HCoV-OC43 spike (Figure 5A & 5D) and HCoV-OC43 nucleocapsid (Figure 5B & 5E) peptide pools even when the individuals with possible prior asymptomatic infection were excluded from the analysis. We next examined cellular responses to non-structural proteins nsp12 and nsp13 (viral RNA dependent RNA polymerase and viral helicase respectively) because these have been associated with abortive SARS-CoV-2 after exposure.^38^ We generated an HCoV-OC43 nsp12/nsp13 peptide pool based on previously described SARS-CoV-2 nsp12 and nsp13 reactive T cell epitopes.^19,27,39–44^ Based on the Immune Epitope Database and Analysis Resource (IEDB) population coverage analysis tool, this combined set of epitopes with their known HLA interactions potentially covers 98.05% (Class I) and 99.3% (Class II) of the world population, and averages 7.82 (Class I) and 5.61 (Class II) epitopes per HLA combination.^45^ The defined SARS-CoV-2 epitopes were mapped to the HCoV-OC43 protein sequence, and HCoV-OC43 nsp12/nsp13 peptides were synthesized and pooled together (Table S4). HCoV-OC43 nsp12/nsp13 reactive CD8^+^ (Figure 5F), but not CD4^+^ T (Figure 5C) cells, were more frequent in individuals with a previous SARS-CoV-2 infection compared to individuals without a known history of SARS-CoV-2 antigen exposure (p = 0.0011, p = 0.0005 after excluding the one with possible “misclassification”) or with a prior COVID-19 vaccination only (p = 0.0148, p = 0.0043 without the seven with possible prior occult infection). Individuals with prior SARS-CoV-2 infection had significantly higher CD8^+^ T cell responses to HCoV-OC43 nsp12/nsp13 peptide pools as compared to a combined group containing those with COVID-19 vaccination and no antigen exposure (p = 0.0076, p = 0.0002 when excluding the eight with possible prior infection). Previous SARS-CoV-2 infection associated with 0.0714 higher percent of nsp12/nsp13 responsive CD8 T cells in multivariable linear regression analysis that accounted for baseline demographics and co-morbidities (Table S5). In this multi-variate model, age greater than 50 years was not a significant predictor for HCoV-OC43 nsp12/nsp13 CD8^+^ T cell responses (Table S5). Importantly, no differences were observed between the groups in the CD4^+^ or CD8^+^ T cell responses to control peptide pools, human cytomegalovirus (CMV) pp65 or CMV, Epstein-Barr virus, and influenza (CEF) (Figure S2). Similar results were observed when stimulation index (SI) was used to calculate the relative T cell activation compared to the negative control condition (Figure S3). This suggests the individuals in the groups did not have baseline differences in cellular reactivity.

**Figure 5.**
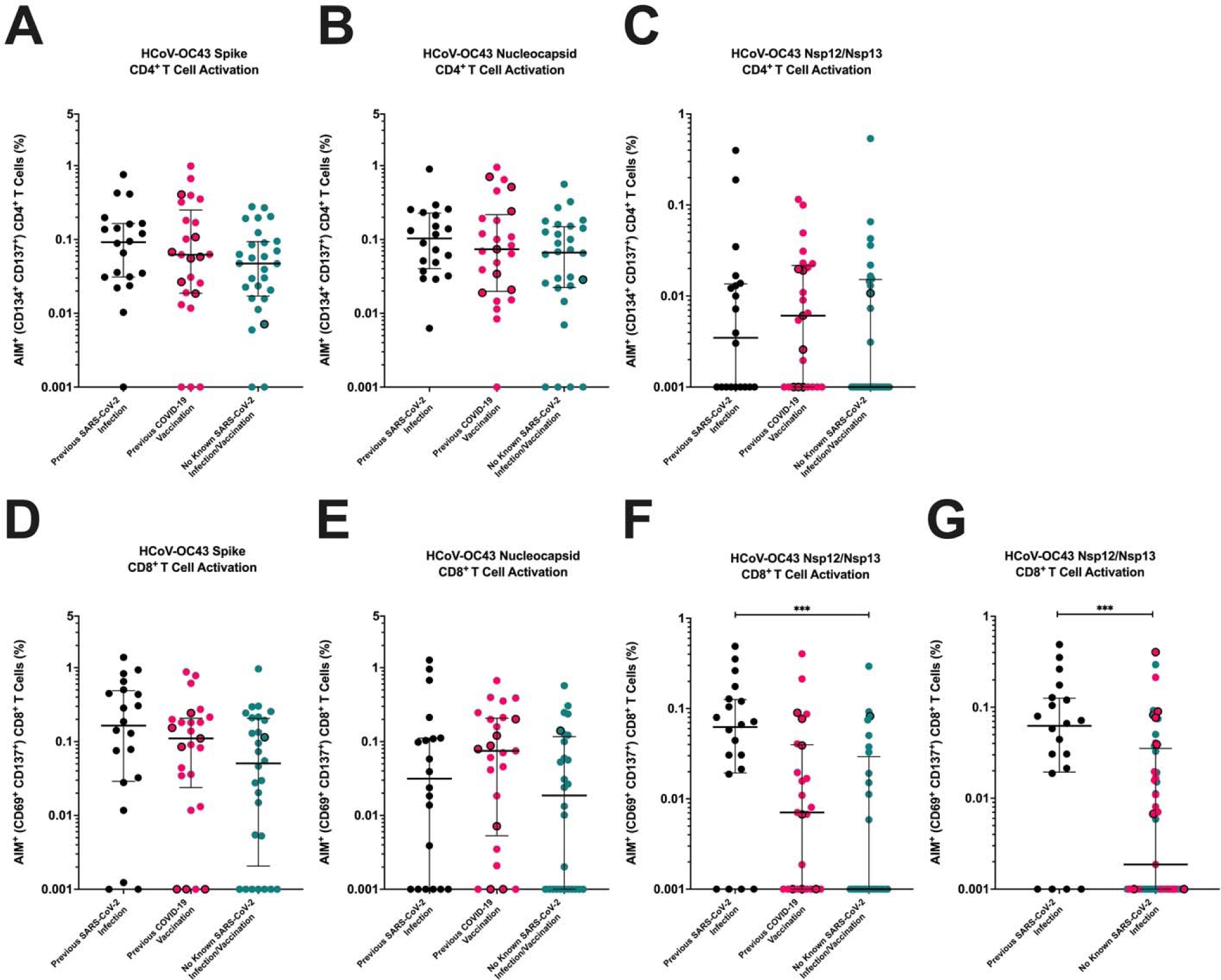
T cell responses to CoV-derived peptide pools among those with different SARS-CoV-2 antigen exposure. Percent of activated CD4^+^ (A-C) and CD8^+^ (D-F) T cells in response to various CoV peptide pools in those with no prior SARS-CoV-2 exposure (teal), prior COVID-19 vaccination but no SARS-CoV-2 infection (pink), and prior documented SARS-CoV-2 infection (black). Black borders represent the eight individuals identified as potentially having prior undocumented or asymptomatic SARS-CoV-2 infection. Patient cells were stimulated with HCoV-OC43 spike (A, D), HCoV-OC43 nucleocapsid (B, E), or HCoV-OC43 nsp12/nsp13 (C, F) peptide pools and activation were measured on CD4^+^ (CD134^+^ CD137^+^) and CD8^+^ (CD69^+^ CD137^+^) T cells by flow cytometry. (G) The same respective data as in panel F, but the individuals with either a previous COVID-19 vaccination or those with known history of SARS-CoV-2 were placed in the same group. Data were background subtracted against the negative control (DMSO only). The dark horizontal lines in each scatter dot plot denote the median and interquartile range. Note, the y-axis varies among the different panels. *** represents p-values <0.001.

## Discussion

Some, but not all, prior studies have implied that previous eCoV immunity can provide protection against SARS-CoV-2 infection and COVID-19 morbidity.^27,46–48^ At BMC, which was the focus for the epidemiological examination in this study, we have previously shown that recent documented eCoV infection ameliorates subsequent COVID-19 disease severity and morbidity, although it likely does not prevent SARS-CoV-2 incidence.^26^ The biological mechanism for this observation remains uncertain although numerous groups have suggested that prior eCoV infection can generate heterotypic humoral and cellular responses against SARS-CoV-2.^25,49^ To our knowledge however, no prior human study has examined if SARS-CoV-2 infection or COVID-19 vaccination protects against a subsequent eCoV infection. Although nearly all adults have preexisting immune responses against the eCoVs, these “common cold” CoV share less genetic similarity with SARS-CoV-2 as compared to the relatedness among different SARS-CoV-2 lineages. Thus, eCoVs can stand as a surrogate for a possible novel heterologous CoV. The incidence of eCoVs after SARS-CoV-2 infection and COVID-19 vaccination provide insights into the prospects for preventing disease from a future novel CoV outbreak, and the immunological characterization can inform future CoV vaccine design.

We find that individuals with a prior SARS-CoV-2 infection, but not COVID-19 vaccination alone, have lower likelihood of having a subsequent symptomatic eCoV infection. Statistically significant differences were only observed for the alpha-CoVs although beta-CoVs incidence was also lower. Alpha-as compared to the beta-CoVs are more distantly related to SARS-CoV-2.^50^ This suggests that SARS-CoV-2 infection as compared to COVID-19 vaccination may provide greater protection against novel genetically disparate CoVs. Similar to our results, SARS-CoV-2 spike immunization alone provided minimal protection against subsequent HCoV-OC43 infection in mice.^51^ In some respects, these animal results support our observations because COVID-19 vaccines predominantly only contain the SARS-CoV-2 spike antigen and no other viral protein. Along with spike directed immunity, individuals with prior SARS-CoV-2 infection harbor additional immune responses against non-spike viral proteins, such as nucleocapsid, nsp12, and nsp13. The non-spike proteins are more conserved among the different CoVs, and thus the immune responses are likely cross-reactive.^52^ The SARS-CoV-2 infection and COVID-19 vaccine generated heterotypic immunity, especially nAbs, may not be able to fully prevent a heterologous CoV infection. After initial infection, however, SARS-CoV-2 heterotypic non-spike cellular responses could lower virus levels and reduce the onset of symptomatic disease.^53,54^ It remains uncertain if SARS-CoV-2 infection also increases the frequency of abortive eCoV infections as compared to those with COVID-19 vaccination alone.

There are several strengths with our observational retrospective results. The data incorporates a relatively large number of individuals even though it was acquired from a single medical center. The large population size allowed us to adjust for various demographics that are likely important for symptomatic eCoVs, such as age, gender, and pre-existing morbidities.^3,55^ Prior SARS-CoV-2 infection associated with lower subsequent eCoV incidence with all the various different statistical methods used to analyze the data. Importantly, in these models, non-eCoV incidence was not different among those with prior SARS-CoV-2 infection and no SARS-CoV-2 antigen experience. This supports the notion that behaviors and unmeasured confounders important for exposure to respiratory infections were likely similar in these groups. On the other hand, COVID-19 vaccinated individuals had greater eCoV infections despite older age associating with lower eCoV infections. This likely reflects differences in exposure to respiratory pathogens as implied by the non-CoV incidence, although it is possible that older individuals have greater immunity from years of prior exposure to the circulating eCoVs. The impact of SARS-CoV-2 infection as compared to COVID-19 vaccination on eCoV incidence should be examined in other cohorts also, although the widespread and often undocumented infection with SARS-CoV-2 Omicron will make a prospective analysis extremely challenging now.^56,57^

We conducted ex-vivo analyses to identify immune mechanisms for the observed protection against symptomatic eCoV infections. These *ex-vivo* studies were on individuals independent from the retrospective analyses with more strict definitions for the different groups. Furthermore, we conducted numerous tests to identify misclassified individuals because it can be difficult to decipher previous asymptomatic and undocumented SARS-CoV-2 infections using single tests alone. We used six different independent measurements to identify misclassified individuals. In general, our methodology revealed similar results as other T cell based methods to differentiate routes of SARS-CoV-2 spike exposure.^34^ It should be noted, however, that no assay or methodology can identify prior SARS-CoV-2 infection with complete certainty.^58^

Numerous previous studies have reported on the importance of nAbs in preventing CoV infections and ameliorating disease severity.^35,36^ Indeed, nAbs are deemed as a correlate of protection against SARS-CoV-2, and as a corollary against CoVs in general.^59^ Animal models also suggest that cross-reactive nAbs can protect against heterologous CoV challenge if the two CoVs have high levels of genetic conservation.^51^ In our analysis, we found that nAbs against SARS-CoV-2, HCoV-OC43, and HCoV-229E were not significantly different among those with prior SARS-CoV-2 infection or COVID-19 vaccination alone. Previous reports have suggested SARS-CoV-2 spike exposure boosts antibody titers against the beta-CoVs, such as HCoV-OC43, and not necessarily against the alpha-CoVs, such as HCoV-229E.^60^ We also found that binding antibodies against HCoV-OC43 spike were increased after SARS-CoV-2 infection, though we did not observe a boost in HCoV-OC43 antibodies in our COVID-19 vaccinated group. In contrast, we observed that HCoV-OC43 neutralization was generally lowest in those with prior documented SARS-CoV-2 infection or COVID-19 vaccination as compared to those without any known SARS-CoV-2 antigen exposure. The lower HCoV-OC43 neutralizing to total antibody binding ratio in those with prior SARS-CoV-2 spike exposure suggests that SARS-CoV-2 spike preferentially induces antibodies that are not critical for HCoV-OC43 neutralization.^61^ It is also possible that the frequency of recent undocumented eCoV, especially HCoV-OC43, infections prior to blood collection was different in the three groups. Similar HCoV-OC43 nucleocapsid directed cellular responses, however, argues against this possibility. In aggregate, our observations suggest that the SARS-CoV-2 infection boosted cross-reactive eCoV binding antibodies are not protective. It should be noted that we did not assess mucosal antibody levels or Fc mediated antibody functions against HCoV-OC43 or HCoV-229E among the groups, and differences in these parameters may be important for protection against symptomatic eCoVs.^62^ Collectively, our studies suggest that pre-existing or de-novo generated plasma nAbs titers against spike do not associate with the lower incidence of symptomatic eCoVs after SARS-CoV-2 infection.

In contrast to nAbs, our ex-vivo investigations imply that cellular responses are important in lowering the incidence of symptomatic eCoVs. The spike-based T cell immunity was similar between those individuals with either a previous SARS-CoV-2 infection and a previous COVID-19 vaccination, which agrees with previous work in the field.^63,64^ In general, the T cell responses against HCoV-OC43 structural proteins were relatively common in all individuals, highlighting the ubiquity of prior eCoV infections.^18^ The primary difference between the individuals with a previous SARS-CoV-2 infection and those with a COVID-19 vaccination were in the elevated CD8^+^ T cell responses to the non-structural antigens, nsp12 and nsp13. Memory T cell responses to these non-structural antigens have been implicated in the protection against SARS-CoV-2 infections.^27,38^ Animal models have also implied that cellular responses against non-structural proteins are important for protection against infection and disease severity.^54,65^ Importantly, the non-structural regions of the different CoVs are generally more conserved and thus cross-reactive responses are more likely preserved.^52,66^ The heterotypic boost of these cross-reactive T cells likely contributes to the killing of CoV infected cells and the clearance of the infection before the onset of symptomatic disease. In our ex-vivo analysis, greater than 50 years of age did not associate with higher HCoV-OC43 nsp12 and nsp13 directed CD8^+^ T cell responses (Table S5). This further argues that the protection against symptomatic eCoV observed in those greater than 50 years of age in the retrospective analysis (Fig. 1) potentially reflects lower exposure to respiratory pathogens rather than enhanced immunity from a greater number of prior HCoV-OC43 infections with increasing age.^67^

There are limitations to our conclusions from the *ex-vivo* analyses. First, we chose to examine spike, nucleocapsid, and nsp12/nsp13 cellular responses only. Cellular responses against other structural and non-structural proteins are likely also important. Cell quantities limited the breadth of the cellular responses that we could examine, and we focused on those that have been deemed most important for disease prevention.^38,68^ We also did not incorporate COVID-19 disease parameters for the individuals with prior SARS-CoV-2 infection in our analyses. The magnitude of the immune responses differ based on COVID-19 disease severity,^69^ and associations between eCoV incidence and mild versus severe COVID-19 disease may further highlight the potentially protective immune responses. It should be noted that our results demonstrate associations and do not prove causation. Documenting lower eCoV incidence among those with as compared to without pre-existing cellular responses against non-structural CoV proteins generated via vaccination or infection will be required to conclude biological mechanisms. These types of human studies will be exceedingly difficult because of the relatively low incidence of symptomatic clinically recognized eCoV infections.

Based on our findings, SARS-CoV-2 infection provides better protection against the heterologous eCoVs, primarily the more genetically distant alpha-eCoVs, compared to spike-based COVID-19 vaccination alone. Even though COVID-19 acute disease severity has been greatly reduced with the widespread institution of COVID-19 vaccination and immunity from prior SARS-CoV-2 infections, our results should not be used to promote SARS-CoV-2 infections as a means to limit future heterologous CoV disease.^70^ SARS-CoV-2 infections continue to pose a significant risk for developing long-term symptoms, termed Long COVID or post-acute sequelae of SARS-CoV-2 infection (PASC).^71,72^ Most next universal CoV vaccines are focused on either eliciting broadly nAbs or inducing T cell responses that target conserved domains in structural proteins shared among diverse CoVs.^73^ While these efforts should continue, we propose that the addition of non-structural antigens to future universal CoV vaccines may induce a more diverse T cell repertoire and recapitulate the heterotypic immune benefit of a natural infection.

## Acknowledgments

We thank the study participants for providing information and donating specimens. We thank the Boston University Flow Cytometry Core and phlebotomy team for their help. The content is solely the responsibility of the authors and does not necessarily represent the official views of the National Institutes of Health.

## Author contributions

MS conceived and designed the study. DJB conducted the experiments. JM and MS collected and tabulated the EMR data. DJB and YML isolated plasma and cells from donated samples. EB, YS, ZS, SC, and DW conducted the Simoa studies. DJB, JW and MS analyzed the data and conducted statistical analyses. DJB and MS wrote the manuscript with editorial assistance from the co-authors.

## Methods

### Participants and Data Collection

We performed a retrospective analysis of the incidence of eCoV and non-eCoV infections among individuals classified into three groups: 1) previous documented SARS-CoV-2 infection; 2) COVID-19 vaccination and no documented or known SARS-CoV-2 infection; and 3) neither exposure. We collected all available comprehensive respiratory panel polymerase chain-reaction (CRP-PCR, *BioFire Diagnostics*) results in the BMC electronic medical records (EMR) performed from November 30, 2020, to October 8, 2021. All individuals less than 18 years of age were excluded from analyses. The start date was chosen because the first eCoV infection at BMC for the 2020 fall/winter season was documented on November 30, 2020. The end date of October 8, 2021 was chosen because this was before the first documented SARS-CoV-2 Omicron infection at BMC. The less symptomatic and widely spread Omicron infections made it difficult to reliably differentiate individuals into the three pre-specified groups.^57^ Individuals with a positive test for HCoV-229E, HCoV-HKU1, HCoV-NL63, or HCoV-OC43 were classified as having an eCoV infection. Each individual only contributed a single data point although they may have had multiple CRP-PCR results during the study period. The date of the positive eCoV infection was used in the analysis for an individual even if they had other CRP-PCR results during the study period. Incidence of non-CoVs served as a control to assess for differences in susceptibility to respiratory infections among the three groups. The date of the first positive non-CoV result was used in the analysis although the individual may have had other pathogens, besides eCoVs, in prior or subsequent CRP-PCRs. Among individuals with multiple CRP-PCRs that did not detect eCoVs or non-CoVs, the date of the last test was used in the analyses. Demographic and clinical information, including prior SARS-CoV-2 vaccination and a previous documented positive SARS-CoV-2 test result, were obtained for these individuals from the EMR.

Blood samples were obtained from BMC patients during a non-COVID-19 related medical visit. Prior to sample collection, all individuals’ status regarding COVID-19 vaccination and prior SARS-CoV-2 infection was confirmed during the consent process. Inclusion criteria included age greater than 18 years of age and medical visit for non-COVID-19 diagnosis. Exclusion included a SARS-CoV-2 positive nasal swab within past 14 days or use of an immunosuppressive drug. Plasma was obtained after centrifugation, and peripheral blood mononuclear cells (PBMCs) were isolated using Ficoll Hypaque density centrifugation methods.

### SARS-CoV-2 Specific IgG Detection

Anti-SARS-CoV-2 IgG antibody levels were assessed using a previously described ultra-sensitive Single Molecule Array (Simoa) multiplexed assay that simultaneously measures anti-SARS-CoV-2 IgG antibodies against receptor binding domain (RBD), and nucleocapsid protein.^30^ Plasma samples were spun down at 4°C for 10 minutes at 2000 x g. The supernatant was then diluted 1000-fold dilution in Homebrew Sample Diluent (Quanterix) before running the assay.

### Pseudovirus Production

Coronavirus-spike expressing pseudoviruses were prepared as previously described.^74^ Briefly, HCoV-229E-spike (SinoBiological, VG40605-CF), HCoV-OC43-spike (SinoBiological, VG40607-CF), or SARS-CoV-2-spike (BEI Resources, NR52310) expression plasmids were transfected into HEK-293T cells. After 24 hours, the transfected cells were infected with VSV-G pseudotyped virus (G*ΔG-VSV) containing the firefly luciferase expression cassette. The pseudovirus-containing cell supernatant was collected after an additional 24 hours, filtered, concentrated, and stored at -80°C.

### Neutralization Assays

Pseudovirus neutralization assays were performed as previously described.^75^ Briefly, plasma was heat inactivated at 56°C for 1 hour, and pseudovirus was incubated in five two-fold serial dilutions, starting with a 1:40 highest dilution, at 37°C. After 1 hour, approximately 1.25×10^4^ Vero E6 cells were added to each well. After 48 hours, luciferase expression was measured by the Promega Bright-Glo Luciferase Assay System (ThermoScientific). Percent neutralization was calculated in comparison to levels of luciferase expression in infected wells without patient plasma. Area under the curve (AUC) was calculated from the curve generated from the neutralizations across the serially diluted plasma.^76^ All neutralizations were tested in triplicate a minimum of two independent times.

### ECoV Spike Specific IgG Detection

Relative HCoV-OC43 and HCoV-229E spike specific IgG levels were detected by ELISA as previously described.^77^ Briefly, a 96-well plate was coated with 2 ug/ml of either HCoV-OC43-spike (SinoBiological, 40607-V08B) or HCoV-229E-spike (SinoBiological, 40605-V08B) antigen for 1 hour at room temperature. Wells were washed thrice with PBS and blocked with casein blocking buffer (Thermo Fisher Scientific, 37528) for 90 minutes. The wells were then washed thrice with PBS, before the wells were incubated with 3-fold dilutions of plasma for 1 hour. Wells were subsequently washed thrice with PBS containing 0.05% Tween 20 (PBST), followed by the addition of anti-human horseradish peroxidase (HRP)-conjugated secondary antibodies for IgG detection (diluted 1:50000, Sigma-Aldrich, A0170) to each well for 30 minutes. The wells were washed with PBST four times, and then 3,3’,5,5’ Tetramethylbenzidine (TMB)-ELISA substrate solution (Thermo Fisher Scientific, 34029) was added and incubated in the dark for 15-20 minutes. The reaction was stopped by the addition of 2M sulfuric acid. Absorbance was measured at 450 nm and the optical density (OD) from the no antigen negative control wells was subtracted from all readings. A standard curve from serial dilutions of the positive control standard (CR3022 IgG, Abcam, 273073) against SARS-CoV-2 receptor binding domain (RBD) (SinoBiological 40592-V08H) was run on each plate. The CR3022 standard curve used to calculate titers (relative units) for each sample by interpolating a four parameter logistic (4PL) curve.

### Peptide Pools

All peptides for a given antigen were reconstituted in dimethyl sulfoxide (DMSO), pooled at equal concentrations, and diluted in phosphate buffered saline (PBS). Aliquots of peptide pools were stored at -80°C. The following peptides pools consisted of overlapping peptides ranging from 13 to 18 amino acids with 10 to 12 amino acids of overlap covering the entire protein region of interest: SARS-CoV-2 spike (n=181, BEI Resources, NR-52402); SARS-CoV-2 nucleocapsid (n=59, BEI Resources, NR-52404); HCoV-OC43 spike (n=226, BEI Resources, NR-53728); HCoV-OC43 nucleocapsid (n=110, JPT, PM-OC43-NCAP-1); and human cytomegalovirus (CMV) pp65 (n=138, NIH HIV Reagent Program, ARP-11549). The peptide pool containing CMV, Epstein-Barr virus, and influenza (CEF) peptides consisted of 32 peptides of 8-11 amino acids covering immunodominant CD8^+^ T cell epitopes (NIH HIV Reagent Program, ARP-9808).

The HCoV-OC43 nsp12/nsp13 peptide pool consisted of 29 peptides of 14 to 16 amino acids in length (Table S4). The defined SARS-CoV-2 epitopes were mapped to the HCoV-OC43 protein sequence (strain ATCC VR-759, NC_006213). Additional amino acids, corresponding to the HCoV-OC43 sequence, were added to the ends of each epitope so each peptide was 14 to 16 amino acids in length. The HCoV-OC43 nsp12/nsp13 peptides were synthesized at greater than 95% purity (GenScript).

### Activation-Induced Marker (AIM) Assay

The AIM assay was performed as described previously.^78,79^ Briefly, PBMC were thawed, washed in Roswell Park Memorial Institute 1640 Medium (RPMI), re-suspended in complete RPMI media with 5% human serum, and rested overnight. The cells were plated at concentration of 1×10^6^ cells/well in a 96-well round bottom plate. For each stimulation, peptides were added to the well at a final concentration of 1 µg/mL, containing less than 0.1% DMSO. Peptide pools containing CEF or CMV pp65 were used as positive controls and media with 0.1% DMSO was used as a negative control. After 24 hours, the cell supernatant was removed, and the cells were collected for additional analysis. Stimulation experiments were performed twice at independent times.

### Cell Staining and Flow Cytometry

Cells from the stimulation experiment were washed twice in PBS and then stained for live/dead cell marker (1:200 dilution, ThermoFisher, L23105) on ice for 20 minutes. Cells were washed in fluorescence-activated cell sorting (FACS) buffer (PBS containing 2% FBS and 2 mM EDTA) and then incubated with human Fc receptor block (1:100 dilution, BioLegend, 422302) for 15 minutes on ice. Cells were then stained with the following antibodies on ice for 30 minutes: Alexa Fluor 647 anti-human CD3 (1:100 dilution, BioLegend, 300321), Alexa Fluor 488 anti-human CD4 (1:200 dilution, BioLegend, 344618), APC/Fire™ 750 anti-human CD8a (1:50 dilution, BioLegend, 300931), PE/Cyanine7 anti-human CD69 (1:50 dilution, BioLegend, 310911), Brilliant Violet 421 anti-human CD134 (OX40) (1:25 dilution, BioLegend, 356147), PE anti-human CD137 (4-1BB (1:100 dilution, BioLegend, 309803). After staining, the cells were washed twice in FACS buffer and samples were analyzed on a BD LSR II Flow Cytometer (BD Biosciences). The resulting flow cytometry data was analyzed using FlowJo software.

### Statistical Analysis

Individuals were initially classified into three groups: 1) those with a prior documented SARS-CoV-2 infection; 2) those with no prior documented SARS-CoV-2 infection and with at least one prior COVID-19 vaccine dose; and 3) those with no prior SARS-CoV-2 infection or COVID-19 vaccination. Vaccinated individuals were deemed fully vaccinated if they had received at least two doses of the Pfizer BioNTech BNT162b2 or Moderna mRNA-1273 COVID-19 vaccine or one dose of the Janssen / Johnson & Johnson Ad26.COV2.S COVID-19 vaccine a minimum of 14 days prior to the CRP-PCR test. No individual had received a COVID-19 vaccine booster. Data are presented as median (interquartile range) for continuous variables and number (percentage) for categorical variables. Comparisons were done using Chi-square for categorical variables and the Kruskal-Wallis or Mann-Whitney test for continuous variables. A Kaplan Meier (KM) curve was used to visually display time to CoV or non-CoV for the three groups. Cox proportional hazards models were developed examining time to either eCoV or non-CoV infection using SARS-CoV-2 infection and COVID-19 vaccination as time-varying covariates. In the KM and Cox proportional hazard models, 11/30/2020 was used as the index or “start” date if SARS-CoV-2 infection or COVID-19 vaccination occurred before 11/30/2020. The start date was the day of exposure if SARS-CoV-2 infection or first COVID-19 vaccination was after 11/30/2020. The event day was the abnormal test or the last follow-up date.

Spike and nucleocapsid indexes were generated by summing log normalized (Log_2_(result + 1)) spike and nucleocapsid SIMOA IgG, AIM-CD4^+^, and AIM-CD8^+^ measurements respectively. In the multivariable linear regression models, antibody, or T cell responses were the dependent variables. The independent predictors varied in the models but included the group categorization or a combined classification based on SARS-CoV-2 spike exposure (SARS-CoV-2 infection and COVID-19 vaccination group) or no known SARS-CoV-2 infection (COVID-19 vaccination and no SARS-CoV-2 antigen experience group). In the multivariable models, covariates included the demographic variables and presence or absence of various comorbidities (Table S1 & S2). All covariates in univariate models significant around the 0.15 level were initially included in multivariable models. Covariates where significance levels fell below 0.1 in multivariable models were then removed. A multiplicative interaction term between SARS-CoV-2 infection and COVID-19 vaccination was examined in Cox proportional hazards models but then removed due to lack of statistical significance. Statistical analyses were performed using GraphPad Prism 8.2.1 and SAS v9.4. A two-sided p-value less than 0.05 was considered statistically significant.

### Study approval

The Institutional Review Board of Boston Medical Center approved this study.

## Data availability

All raw data will be made available from the corresponding author upon request and signature of data transfer agreement.

## Supplemental Information Titles and Legends

**Figure S1. AIM assay gating strategy and analysis.** Related to Figure 4. (A) Representative charts depicting the flow cytometry gating strategy for defining CD4^+^ and CD8^+^ T cell populations. (B) Examples of the flow cytometry gating strategy for measuring antigen specific CD4^+^ (CD134^+^ CD137^+^) and CD8^+^ (CD69^+^ CD137^+^) T cells after stimulation with control or experimental conditions. Of note, this individual had received a COVID-19 vaccination, but had no history of a SARS-CoV-2 infection.

**Figure S2. T cell responses to CMV and CEF peptide pools among those with different SARS-CoV-2 antigen exposure.** Percent of activated CD4^+^ (A, B) and CD8^+^ T cells (C, D) in response to human cytomegalovirus (CMV) pp65 (A, C) or CMV, Epstein-Barr virus, and influenza (CEF) (B, D) peptide pools in those with no prior SARS-CoV-2 exposure (teal), prior COVID-19 vaccination but no SARS-CoV-2 infection (pink), and prior documented SARS-CoV-2 infection (black). Black borders represent the eight individuals identified as potentially having prior undocumented or asymptomatic SARS-CoV-2 infection. Data were background subtracted against the negative control (DMSO only). Note, the y-axis varies among the different panels.

**Figure S3. Alternative analysis of T cell responses to CoV-derived peptide pools among those with different SARS-CoV-2 antigen exposure.** A calculation of relative CD4^+^ (top row) and CD8^+^ T (bottom row) cell responses to various CoV peptide pools in those with no prior SARS-CoV-2 exposure (teal), prior COVID-19 vaccination but no SARS-CoV-2 infection (pink), and prior documented SARS-CoV-2 infection (black). Black borders represent the eight individuals identified as potentially having prior undocumented or asymptomatic SARS-CoV-2 infection. The data is the same as in Figures 3 and 5, but the percent of activated T cells for each experimental condition was divided by the negative control (DMSO only) condition to calculate a stimulation index. The dark horizontal lines in each scatter dot plot denote the median and interquartile range. Note, the y-axis varies among the different panels. *, **, ***, **** represent p-values <0.05, <0.01, <0.001, <0.0001 respectively.

**Table S1. Demographics of the three groups in the retrospective cohort analyses.**

**Table S2. Demographics of the individuals with collected blood specimens.**

**Table S3: Multi-variable linear regression analysis for predictors of HCoV-OC43 spike binding antibodies.**

**Table S4. HCoV-OC43 nsp12/nsp12 peptides.**

**Table S5: Multi-variable linear regression analysis for predictors of HCoV-OC43 nsp12/nsp13 CD8+ T cell responses.**

## References

1. V’kovski, P., Kratzel, A., Steiner, S., Stalder, H., and Thiel, V. (2021). Coronavirus biology and replication: implications for SARS-CoV-2. Nat. Rev. Microbiol. 19, 155–170. 10.1038/s41579-020-00468-6.

2. Gaunt, E.R., Hardie, A., Claas, E.C.J., Simmonds, P., and Templeton, K.E. (2010). Epidemiology and clinical presentations of the four human coronaviruses 229E, HKU1, NL63, and OC43 detected over 3 years using a novel multiplex real-time PCR method. J. Clin. Microbiol. 48, 2940–2947. 10.1128/JCM.00636-10.

3. Walsh, E.E., Shin, J.H., and Falsey, A.R. (2013). Clinical impact of human coronaviruses 229E and OC43 infection in diverse adult populations. J. Infect. Dis. 208, 1634–1642. 10.1093/infdis/jit393.

4. Fung, T.S., and Liu, D.X. (2021). Similarities and Dissimilarities of COVID-19 and Other Coronavirus Diseases. Annu. Rev. Microbiol. 75, 19–47. 10.1146/annurev-micro-110520-023212.

5. Chen, G., Wu, D., Guo, W., Cao, Y., Huang, D., Wang, H., Wang, T., Zhang, X., Chen, H., Yu, H., et al. Clinical and immunological features of severe and moderate coronavirus disease 2019. J. Clin. Invest. 130, 2620–2629. 10.1172/JCI137244.

6. Hu, B., Guo, H., Zhou, P., and Shi, Z.-L. (2021). Characteristics of SARS-CoV-2 and COVID-19. Nat. Rev. Microbiol. 19, 141–154. 10.1038/s41579-020-00459-7.

7. Baden, L.R., El Sahly, H.M., Essink, B., Kotloff, K., Frey, S., Novak, R., Diemert, D., Spector, S.A., Rouphael, N., Creech, C.B., et al. (2021). Efficacy and Safety of the mRNA-1273 SARS-CoV-2 Vaccine. N. Engl. J. Med. 384, 403–416. 10.1056/NEJMoa2035389.

8. Polack, F.P., Thomas, S.J., Kitchin, N., Absalon, J., Gurtman, A., Lockhart, S., Perez, J.L., Pérez Marc, G., Moreira, E.D., Zerbini, C., et al. (2020). Safety and Efficacy of the BNT162b2 mRNA Covid-19 Vaccine. N. Engl. J. Med. 383, 2603–2615. 10.1056/NEJMoa2034577.

9. Andrews, N., Tessier, E., Stowe, J., Gower, C., Kirsebom, F., Simmons, R., Gallagher, E., Thelwall, S., Groves, N., Dabrera, G., et al. (2022). Duration of Protection against Mild and Severe Disease by Covid-19 Vaccines. N. Engl. J. Med. 386, 340–350. 10.1056/NEJMoa2115481.

10. Andrews, N., Stowe, J., Kirsebom, F., Toffa, S., Rickeard, T., Gallagher, E., Gower, C., Kall, M., Groves, N., O’Connell, A.-M., et al. (2022). Covid-19 Vaccine Effectiveness against the Omicron (B.1.1.529) Variant. N. Engl. J. Med. 386, 1532–1546. 10.1056/NEJMoa2119451.

11. Lau, J.J., Cheng, S.M.S., Leung, K., Lee, C.K., Hachim, A., Tsang, L.C.H., Yam, K.W.H., Chaothai, S., Kwan, K.K.H., Chai, Z.Y.H., et al. (2023). Real-world COVID-19 vaccine effectiveness against the Omicron BA.2 variant in a SARS-CoV-2 infection-naive population. Nat. Med. 29, 348–357. 10.1038/s41591-023-02219-5.

12. Pajon, R., Doria-Rose, N.A., Shen, X., Schmidt, S.D., O’Dell, S., McDanal, C., Feng, W., Tong, J., Eaton, A., Maglinao, M., et al. (2022). SARS-CoV-2 Omicron Variant Neutralization after mRNA-1273 Booster Vaccination. N. Engl. J. Med. 386, 1088–1091. 10.1056/NEJMc2119912.

13. Miller, J., Hachmann, N.P., Collier, A.Y., Lasrado, N., Mazurek, C.R., Patio, R.C., Powers, O., Surve, N., Theiler, J., Korber, B., et al. (2023). Substantial Neutralization Escape by SARS-CoV-2 Omicron Variants BQ.1.1 and XBB.1. N. Engl. J. Med. 388, 662–664. 10.1056/NEJMc2214314.

14. Rössler, A., Riepler, L., Bante, D., von Laer, D., and Kimpel, J. (2022). SARS-CoV-2 Omicron Variant Neutralization in Serum from Vaccinated and Convalescent Persons. N. Engl. J. Med. 386, 698–700. 10.1056/NEJMc2119236.

15. Tan, A.T., Linster, M., Tan, C.W., Le Bert, N., Chia, W.N., Kunasegaran, K., Zhuang, Y., Tham, C.Y.L., Chia, A., Smith, G.J.D., et al. (2021). Early induction of functional SARS-CoV-2-specific T cells associates with rapid viral clearance and mild disease in COVID-19 patients. Cell Rep. 34, 108728. 10.1016/j.celrep.2021.108728.

16. Bange, E.M., Han, N.A., Wileyto, P., Kim, J.Y., Gouma, S., Robinson, J., Greenplate, A.R., Hwee, M.A., Porterfield, F., Owoyemi, O., et al. (2021). CD8+ T cells contribute to survival in patients with COVID-19 and hematologic cancer. Nat. Med. 27, 1280–1289. 10.1038/s41591-021-01386-7.

17. Le Bert, N., Clapham, H.E., Tan, A.T., Chia, W.N., Tham, C.Y.L., Lim, J.M., Kunasegaran, K., Tan, L.W.L., Dutertre, C.-A., Shankar, N., et al. (2021). Highly functional virus-specific cellular immune response in asymptomatic SARS-CoV-2 infection. J. Exp. Med. 218, e20202617. 10.1084/jem.20202617.

18. Grifoni, A., Weiskopf, D., Ramirez, S.I., Mateus, J., Dan, J.M., Moderbacher, C.R., Rawlings, S.A., Sutherland, A., Premkumar, L., Jadi, R.S., et al. (2020). Targets of T Cell Responses to SARS-CoV-2 Coronavirus in Humans with COVID-19 Disease and Unexposed Individuals. Cell 181, 1489–1501.e15. 10.1016/j.cell.2020.05.015.

19. Tarke, A., Sidney, J., Kidd, C.K., Dan, J.M., Ramirez, S.I., Yu, E.D., Mateus, J., da Silva Antunes, R., Moore, E., Rubiro, P., et al. (2021). Comprehensive analysis of T cell immunodominance and immunoprevalence of SARS-CoV-2 epitopes in COVID-19 cases. Cell Rep. Med. 2, 100204. 10.1016/j.xcrm.2021.100204.

20. Jungreis, I., Sealfon, R., and Kellis, M. (2021). SARS-CoV-2 gene content and COVID-19 mutation impact by comparing 44 Sarbecovirus genomes. Nat. Commun. 12, 2642. 10.1038/s41467-021-22905-7.

21. Carabelli, A.M., Peacock, T.P., Thorne, L.G., Harvey, W.T., Hughes, J., de Silva, T.I., Peacock, S.J., Barclay, W.S., de Silva, T.I., Towers, G.J., et al. (2023). SARS-CoV-2 variant biology: immune escape, transmission and fitness. Nat. Rev. Microbiol. 21, 162–177. 10.1038/s41579-022-00841-7.

22. Feng, C., Shi, J., Fan, Q., Wang, Y., Huang, H., Chen, F., Tang, G., Li, Y., Li, P., Li, J., et al. (2021). Protective humoral and cellular immune responses to SARS-CoV-2 persist up to 1 year after recovery. Nat. Commun. 12, 4984. 10.1038/s41467-021-25312-0.

23. Bilich, T., Nelde, A., Heitmann, J.S., Maringer, Y., Roerden, M., Bauer, J., Rieth, J., Wacker, M., Peter, A., Hörber, S., et al. (2021). T cell and antibody kinetics delineate SARS-CoV-2 peptides mediating long-term immune responses in COVID-19 convalescent individuals. Sci. Transl. Med. 13, eabf7517. 10.1126/scitranslmed.abf7517.

24. Ng, K.W., Faulkner, N., Cornish, G.H., Rosa, A., Harvey, R., Hussain, S., Ulferts, R., Earl, C., Wrobel, A.G., Benton, D.J., et al. (2020). Preexisting and de novo humoral immunity to SARS-CoV-2 in humans. Science 370, 1339–1343. 10.1126/science.abe1107.

25. Le Bert, N., Tan, A.T., Kunasegaran, K., Tham, C.Y.L., Hafezi, M., Chia, A., Chng, M.H.Y., Lin, M., Tan, N., Linster, M., et al. (2020). SARS-CoV-2-specific T cell immunity in cases of COVID-19 and SARS, and uninfected controls. Nature 584, 457–462. 10.1038/s41586-020-2550-z.

26. Sagar, M., Reifler, K., Rossi, M., Miller, N.S., Sinha, P., White, L.F., and Mizgerd, J.P. (2021). Recent endemic coronavirus infection is associated with less-severe COVID-19. J. Clin. Invest. 131, e143380. 10.1172/JCI143380.

27. Kundu, R., Narean, J.S., Wang, L., Fenn, J., Pillay, T., Fernandez, N.D., Conibear, E., Koycheva, A., Davies, M., Tolosa-Wright, M., et al. (2022). Cross-reactive memory T cells associate with protection against SARS-CoV-2 infection in COVID-19 contacts. Nat. Commun. 13, 80. 10.1038/s41467-021-27674-x.

28. Sinha, P., Reifler, K., Rossi, M., and Sagar, M. (2021). Coronavirus Disease 2019 Mitigation Strategies Were Associated With Decreases in Other Respiratory Virus Infections. Open Forum Infect. Dis. 8, ofab105. 10.1093/ofid/ofab105.

29. Kalish, H., Klumpp-Thomas, C., Hunsberger, S., Baus, H.A., Fay, M.P., Siripong, N., Wang, J., Hicks, J., Mehalko, J., Travers, J., et al. (2021). Undiagnosed SARS-CoV-2 seropositivity during the first 6 months of the COVID-19 pandemic in the United States. Sci. Transl. Med. 13, eabh3826. 10.1126/scitranslmed.abh3826.

30. Norman, M., Gilboa, T., Ogata, A.F., Maley, A.M., Cohen, L., Busch, E.L., Lazarovits, R., Mao, C.-P., Cai, Y., Zhang, J., et al. (2020). Ultrasensitive high-resolution profiling of early seroconversion in patients with COVID-19. Nat. Biomed. Eng. 4, 1180–1187. 10.1038/s41551-020-00611-x.

31. Indenbaum, V., Koren, R., Katz-Likvornik, S., Yitzchaki, M., Halpern, O., Regev-Yochay, G., Cohen, C., Biber, A., Feferman, T., Saban, N.C., et al. (2020). Testing IgG antibodies against the RBD of SARS-CoV-2 is sufficient and necessary for COVID-19 diagnosis. PLOS ONE 15, e0241164. 10.1371/journal.pone.0241164.

32. Burbelo, P.D., Riedo, F.X., Morishima, C., Rawlings, S., Smith, D., Das, S., Strich, J.R., Chertow, D.S., Davey, R.T., Jr, and Cohen, J.I. (2020). Sensitivity in Detection of Antibodies to Nucleocapsid and Spike Proteins of Severe Acute Respiratory Syndrome Coronavirus 2 in Patients With Coronavirus Disease 2019. J. Infect. Dis. 222, 206–213. 10.1093/infdis/jiaa273.

33. Hillig, T., Kristensen, J.R., Brasen, C.L., Brandslund, I., Olsen, D.A., Davidsen, C., Madsen, J.S., Jensen, C.A., Hansen, Y.B.L., and Friis-Hansen, L. (2023). Sensitivity and performance of three novel quantitative assays of SARS-CoV-2 nucleoprotein in blood. Sci. Rep. 13, 2868. 10.1038/s41598-023-29973-3.

34. Yu, E.D., Wang, E., Garrigan, E., Goodwin, B., Sutherland, A., Tarke, A., Chang, J., Gálvez, R.I., Mateus, J., Ramirez, S.I., et al. (2022). Development of a T cell-based immunodiagnostic system to effectively distinguish SARS-CoV-2 infection and COVID-19 vaccination status. Cell Host Microbe 30, 388–399.e3. 10.1016/j.chom.2022.02.003.

35. Feng, S., Phillips, D.J., White, T., Sayal, H., Aley, P.K., Bibi, S., Dold, C., Fuskova, M., Gilbert, S.C., Hirsch, I., et al. (2021). Correlates of protection against symptomatic and asymptomatic SARS-CoV-2 infection. Nat. Med. 27, 2032–2040. 10.1038/s41591-021-01540-1.

36. Khoury, D.S., Cromer, D., Reynaldi, A., Schlub, T.E., Wheatley, A.K., Juno, J.A., Subbarao, K., Kent, S.J., Triccas, J.A., and Davenport, M.P. (2021). Neutralizing antibody levels are highly predictive of immune protection from symptomatic SARS-CoV-2 infection. Nat. Med. 27, 1205–1211. 10.1038/s41591-021-01377-8.

37. Cohen, K.W., Linderman, S.L., Moodie, Z., Czartoski, J., Lai, L., Mantus, G., Norwood, C., Nyhoff, L.E., Edara, V.V., Floyd, K., et al. (2021). Longitudinal analysis shows durable and broad immune memory after SARS-CoV-2 infection with persisting antibody responses and memory B and T cells. Cell Rep. Med. 2, 100354. 10.1016/j.xcrm.2021.100354.

38. Swadling, L., Diniz, M.O., Schmidt, N.M., Amin, O.E., Chandran, A., Shaw, E., Pade, C., Gibbons, J.M., Le Bert, N., Tan, A.T., et al. (2021). Pre-existing polymerase-specific T cells expand in abortive seronegative SARS-CoV-2. Nature, 1–8. 10.1038/s41586-021-04186-8.

39. Saini, S.K., Hersby, D.S., Tamhane, T., Povlsen, H.R., Amaya Hernandez, S.P., Nielsen, M., Gang, A.O., and Hadrup, S.R. (2021). SARS-CoV-2 genome-wide T cell epitope mapping reveals immunodominance and substantial CD8+ T cell activation in COVID-19 patients. Sci. Immunol. 6, eabf7550. 10.1126/sciimmunol.abf7550.

40. Kared, H., Redd, A.D., Bloch, E.M., Bonny, T.S., Sumatoh, H., Kairi, F., Carbajo, D., Abel, B., Newell, E.W., Bettinotti, M.P., et al. (2021). SARS-CoV-2-specific CD8+ T cell responses in convalescent COVID-19 individuals. J. Clin. Invest. 131, 145476. 10.1172/JCI145476.

41. Ferretti, A.P., Kula, T., Wang, Y., Nguyen, D.M.V., Weinheimer, A., Dunlap, G.S., Xu, Q., Nabilsi, N., Perullo, C.R., Cristofaro, A.W., et al. (2020). Unbiased Screens Show CD8+ T Cells of COVID-19 Patients Recognize Shared Epitopes in SARS-CoV-2 that Largely Reside outside the Spike Protein. Immunity 53, 1095–1107.e3. 10.1016/j.immuni.2020.10.006.

42. Mateus, J., Grifoni, A., Tarke, A., Sidney, J., Ramirez, S.I., Dan, J.M., Burger, Z.C., Rawlings, S.A., Smith, D.M., Phillips, E., et al. (2020). Selective and cross-reactive SARS-CoV-2 T cell epitopes in unexposed humans. Science 370, 89–94. 10.1126/science.abd3871.

43. Schulien, I., Kemming, J., Oberhardt, V., Wild, K., Seidel, L.M., Killmer, S., Sagar Daul, F., Salvat Lago, M., Decker, A., et al. (2021). Characterization of pre-existing and induced SARS-CoV-2-specific CD8+ T cells. Nat. Med. 27, 78–85. 10.1038/s41591-020-01143-2.

44. Nelde, A., Bilich, T., Heitmann, J.S., Maringer, Y., Salih, H.R., Roerden, M., Lübke, M., Bauer, J., Rieth, J., Wacker, M., et al. (2021). SARS-CoV-2-derived peptides define heterologous and COVID-19-induced T cell recognition. Nat. Immunol. 22, 74–85. 10.1038/s41590-020-00808-x.

45. Bui, H.-H., Sidney, J., Dinh, K., Southwood, S., Newman, M.J., and Sette, A. (2006). Predicting population coverage of T-cell epitope-based diagnostics and vaccines. BMC Bioinformatics 7, 153. 10.1186/1471-2105-7-153.

46. Loyal, L., Braun, J., Henze, L., Kruse, B., Dingeldey, M., Reimer, U., Kern, F., Schwarz, T., Mangold, M., Unger, C., et al. Cross-reactive CD4+ T cells enhance SARS-CoV-2 immune responses upon infection and vaccination. Science 374, eabh1823. 10.1126/science.abh1823.

47. Dugas, M., Grote-Westrick, T., Merle, U., Fontenay, M., Kremer, A.E., Hanses, F., Vollenberg, R., Lorentzen, E., Tiwari-Heckler, S., Duchemin, J., et al. (2021). Lack of antibodies against seasonal coronavirus OC43 nucleocapsid protein identifies patients at risk of critical COVID-19. J. Clin. Virol. 139, 104847. 10.1016/j.jcv.2021.104847.

48. Abela, I.A., Pasin, C., Schwarzmüller, M., Epp, S., Sickmann, M.E., Schanz, M.M., Rusert, P., Weber, J., Schmutz, S., Audigé, A., et al. (2021). Multifactorial seroprofiling dissects the contribution of pre-existing human coronaviruses responses to SARS-CoV-2 immunity. Nat. Commun. 12, 6703. 10.1038/s41467-021-27040-x.

49. Song, G., He, W., Callaghan, S., Anzanello, F., Huang, D., Ricketts, J., Torres, J.L., Beutler, N., Peng, L., Vargas, S., et al. (2021). Cross-reactive serum and memory B-cell responses to spike protein in SARS-CoV-2 and endemic coronavirus infection. Nat. Commun. 12, 2938. 10.1038/s41467-021-23074-3.

50. Chen, Z., Boon, S.S., Wang, M.H., Chan, R.W.Y., and Chan, P.K.S. (2021). Genomic and evolutionary comparison between SARS-CoV-2 and other human coronaviruses. J. Virol. Methods 289, 114032. 10.1016/j.jviromet.2020.114032.

51. Dangi, T., Palacio, N., Sanchez, S., Park, M., Class, J., Visvabharathy, L., Ciucci, T., Koralnik, I.J., Richner, J.M., and Penaloza-MacMaster, P. (2021). Cross-protective immunity following coronavirus vaccination and coronavirus infection. J. Clin. Invest. 131, e151969. 10.1172/JCI151969.

52. Westphal, T., Mader, M., Karsten, H., Cords, L., Knapp, M., Schulte, S., Hermanussen, L., Peine, S., Ditt, V., Grifoni, A., et al. (2023). Evidence for broad cross-reactivity of the SARS-CoV-2 NSP12-directed CD4+ T-cell response with pre-primed responses directed against common cold coronaviruses. Front. Immunol. 14, 1182504. 10.3389/fimmu.2023.1182504.

53. Dangi, T., Class, J., Palacio, N., Richner, J.M., and MacMaster, P.P. (2021). Combining spike- and nucleocapsid-based vaccines improves distal control of SARS-CoV-2. Cell Rep. 36, 109664. 10.1016/j.celrep.2021.109664.

54. Dagotto, G., Ventura, J.D., Martinez, D.R., Anioke, T., Chung, B.S., Siamatu, M., Barrett, J., Miller, J., Schäfer, A., Yu, J., et al. (2022). Immunogenicity and protective efficacy of a rhesus adenoviral vaccine targeting conserved COVID-19 replication transcription complex. NPJ Vaccines 7, 125. 10.1038/s41541-022-00553-2.

55. Cummings, D.A.T., Radonovich, L.J., Gorse, G.J., Gaydos, C.A., Bessesen, M.T., Brown, A.C., Gibert, C.L., Hitchings, M.D.T., Lessler, J., Nyquist, A.-C., et al. (2021). Risk Factors for Healthcare Personnel Infection With Endemic Coronaviruses (HKU1, OC43, NL63, 229E): Results from the Respiratory Protection Effectiveness Clinical Trial (ResPECT). Clin. Infect. Dis. Off. Publ. Infect. Dis. Soc. Am. 73, e4428–e4432. 10.1093/cid/ciaa900.

56. Baker, J.M., Nakayama, J.Y., O’Hegarty, M., McGowan, A., Teran, R.A., Bart, S.M., Mosack, K., Roberts, N., Campos, B., Paegle, A., et al. (2022). SARS-CoV-2 B.1.1.529 (Omicron) Variant Transmission Within Households - Four U.S. Jurisdictions, November 2021-February 2022. MMWR Morb. Mortal. Wkly. Rep. 71, 341–346. 10.15585/mmwr.mm7109e1.

57. Clarke, K.E.N., Jones, J.M., Deng, Y., Nycz, E., Lee, A., Iachan, R., Gundlapalli, A.V., Hall, A.J., and MacNeil, A. (2022). Seroprevalence of Infection-Induced SARS-CoV-2 Antibodies - United States, September 2021-February 2022. MMWR Morb. Mortal. Wkly. Rep. 71, 606–608. 10.15585/mmwr.mm7117e3.

58. Loesche, M., Karlson, E.W., Talabi, O., Zhou, G., Boutin, N., Atchley, R., Loevinsohn, G., Chang, J.B.P., Hasdianda, M.A., Okenla, A., et al. Longitudinal SARS-CoV-2 Nucleocapsid Antibody Kinetics, Seroreversion, and Implications for Seroepidemiologic Studies. Emerg. Infect. Dis. 28, 1859–1862. 10.3201/eid2809.220729.

59. Huang, A.T., Garcia-Carreras, B., Hitchings, M.D.T., Yang, B., Katzelnick, L.C., Rattigan, S.M., Borgert, B.A., Moreno, C.A., Solomon, B.D., Trimmer-Smith, L., et al. (2020). A systematic review of antibody mediated immunity to coronaviruses: kinetics, correlates of protection, and association with severity. Nat. Commun. 11, 4704. 10.1038/s41467-020-18450-4.

60. Anderson, E.M., Goodwin, E.C., Verma, A., Arevalo, C.P., Bolton, M.J., Weirick, M.E., Gouma, S., McAllister, C.M., Christensen, S.R., Weaver, J., et al. (2021). Seasonal human coronavirus antibodies are boosted upon SARS-CoV-2 infection but not associated with protection. Cell 184, 1858–1864.e10. 10.1016/j.cell.2021.02.010.

61. Ng, K.W., Faulkner, N., Finsterbusch, K., Wu, M., Harvey, R., Hussain, S., Greco, M., Liu, Y., Kjaer, S., Swanton, C., et al. (2022). SARS-CoV-2 S2-targeted vaccination elicits broadly neutralizing antibodies. Sci. Transl. Med. 14, eabn3715. 10.1126/scitranslmed.abn3715.

62. Tang, J., Zeng, C., Cox, T.M., Li, C., Son, Y.M., Cheon, I.S., Wu, Y., Behl, S., Taylor, J.J., Chakaraborty, R., et al. (2022). Respiratory mucosal immunity against SARS-CoV-2 after mRNA vaccination. Sci. Immunol. 7, eadd4853. 10.1126/sciimmunol.add4853.

63. Hurme, A., Jalkanen, P., Heroum, J., Liedes, O., Vara, S., Melin, M., Teräsjärvi, J., He, Q., Pöysti, S., Hänninen, A., et al. (2022). Long-Lasting T Cell Responses in BNT162b2 COVID-19 mRNA Vaccinees and COVID-19 Convalescent Patients. Front. Immunol. 13, 869990. 10.3389/fimmu.2022.869990.

64. Naranbhai, V., Nathan, A., Kaseke, C., Berrios, C., Khatri, A., Choi, S., Getz, M.A., Tano-Menka, R., Ofoman, O., Gayton, A., et al. (2022). T cell reactivity to the SARS-CoV-2 Omicron variant is preserved in most but not all individuals. Cell 185, 1041–1051.e6. 10.1016/j.cell.2022.01.029.

65. Tai, W., Feng, S., Chai, B., Lu, S., Zhao, G., Chen, D., Yu, W., Ren, L., Shi, H., Lu, J., et al. (2023). An mRNA-based T-cell-inducing antigen strengthens COVID-19 vaccine against SARS-CoV-2 variants. Nat. Commun. 14, 2962. 10.1038/s41467-023-38751-8.

66. Nesterenko, P.A., McLaughlin, J., Tsai, B.L., Burton Sojo, G., Cheng, D., Zhao, D., Mao, Z., Bangayan, N.J., Obusan, M.B., Su, Y., et al. (2021). HLA-A∗02:01 restricted T cell receptors against the highly conserved SARS-CoV-2 polymerase cross-react with human coronaviruses. Cell Rep. 37, 110167. 10.1016/j.celrep.2021.110167.

67. Edridge, A.W.D., Kaczorowska, J., Hoste, A.C.R., Bakker, M., Klein, M., Loens, K., Jebbink, M.F., Matser, A., Kinsella, C.M., Rueda, P., et al. (2020). Seasonal coronavirus protective immunity is short-lasting. Nat. Med. 26, 1691–1693. 10.1038/s41591-020-1083-1.

68. Hajnik, R.L., Plante, J.A., Liang, Y., Alameh, M.-G., Tang, J., Bonam, S.R., Zhong, C., Adam, A., Scharton, D., Rafael, G.H., et al. (2022). Dual spike and nucleocapsid mRNA vaccination confer protection against SARS-CoV-2 Omicron and Delta variants in preclinical models. Sci. Transl. Med. 14, eabq1945. 10.1126/scitranslmed.abq1945.

69. Dan, J.M., Mateus, J., Kato, Y., Hastie, K.M., Yu, E.D., Faliti, C.E., Grifoni, A., Ramirez, S.I., Haupt, S., Frazier, A., et al. (2021). Immunological memory to SARS-CoV-2 assessed for up to 8 months after infection. Science 371. 10.1126/science.abf4063.

70. Tenforde, M.W., Self, W.H., Adams, K., Gaglani, M., Ginde, A.A., McNeal, T., Ghamande, S., Douin, D.J., Talbot, H.K., Casey, J.D., et al. (2021). Association Between mRNA Vaccination and COVID-19 Hospitalization and Disease Severity. JAMA 326, 2043–2054. 10.1001/jama.2021.19499.

71. Davis, H.E., McCorkell, L., Vogel, J.M., and Topol, E.J. (2023). Long COVID: major findings, mechanisms and recommendations. Nat. Rev. Microbiol. 21, 133–146. 10.1038/s41579-022-00846-2.

72. Groff, D., Sun, A., Ssentongo, A.E., Ba, D.M., Parsons, N., Poudel, G.R., Lekoubou, A., Oh, J.S., Ericson, J.E., Ssentongo, P., et al. (2021). Short-term and Long-term Rates of Postacute Sequelae of SARS-CoV-2 Infection: A Systematic Review. JAMA Netw. Open 4, e2128568. 10.1001/jamanetworkopen.2021.28568.

73. Su, S., Li, W., and Jiang, S. (2022). Developing pan-β-coronavirus vaccines against emerging SARS-CoV-2 variants of concern. Trends Immunol. 43, 170–172. 10.1016/j.it.2022.01.009.

74. Whitt, M.A. (2010). Generation of VSV pseudotypes using recombinant ΔG-VSV for studies on virus entry, identification of entry inhibitors, and immune responses to vaccines. J. Virol. Methods 169, 365–374. 10.1016/j.jviromet.2010.08.006.

75. Nie, J., Li, Q., Wu, J., Zhao, C., Hao, H., Liu, H., Zhang, L., Nie, L., Qin, H., Wang, M., et al. (2020). Quantification of SARS-CoV-2 neutralizing antibody by a pseudotyped virus-based assay. Nat. Protoc. 15, 3699–3715. 10.1038/s41596-020-0394-5.

76. Yu, X., Gilbert, P.B., Hioe, C.E., Zolla-Pazner, S., and Self, S.G. (2012). Statistical approaches to analyzing HIV-1 neutralizing antibody assay data. Stat. Biopharm. Res. 4, 1–13. 10.1080/19466315.2011.633860.

77. Yuen, R.R., Steiner, D., Pihl, R.M.F., Chavez, E., Olson, A., Smith, E.L., Baird, L.A., Korkmaz, F., Urick, P., Sagar, M., et al. (2021). Novel ELISA Protocol Links Pre-Existing SARS-CoV-2 Reactive Antibodies With Endemic Coronavirus Immunity and Age and Reveals Improved Serologic Identification of Acute COVID-19 via Multi-Parameter Detection. Front. Immunol. 12. 10.3389/fimmu.2021.614676.

78. Dan, J.M., Lindestam Arlehamn, C.S., Weiskopf, D., da Silva Antunes, R., Havenar-Daughton, C., Reiss, S.M., Brigger, M., Bothwell, M., Sette, A., and Crotty, S. (2016). A Cytokine-Independent Approach To Identify Antigen-Specific Human Germinal Center T Follicular Helper Cells and Rare Antigen-Specific CD4+ T Cells in Blood. J. Immunol. 197, 983–993. 10.4049/jimmunol.1600318.

79. Reiss, S., Baxter, A.E., Cirelli, K.M., Dan, J.M., Morou, A., Daigneault, A., Brassard, N., Silvestri, G., Routy, J.-P., Havenar-Daughton, C., et al. (2017). Comparative analysis of activation induced marker (AIM) assays for sensitive identification of antigen-specific CD4 T cells. PloS One 12, e0186998. 10.1371/journal.pone.0186998.

