## Supplemental Figures and Tables for "Heterotypic responses against nsp12/nsp13 from prior SARS-CoV-2 infection associates with lower subsequent endemic coronavirus incidence"

### Figure S1. AIM assay gating strategy and analysis.

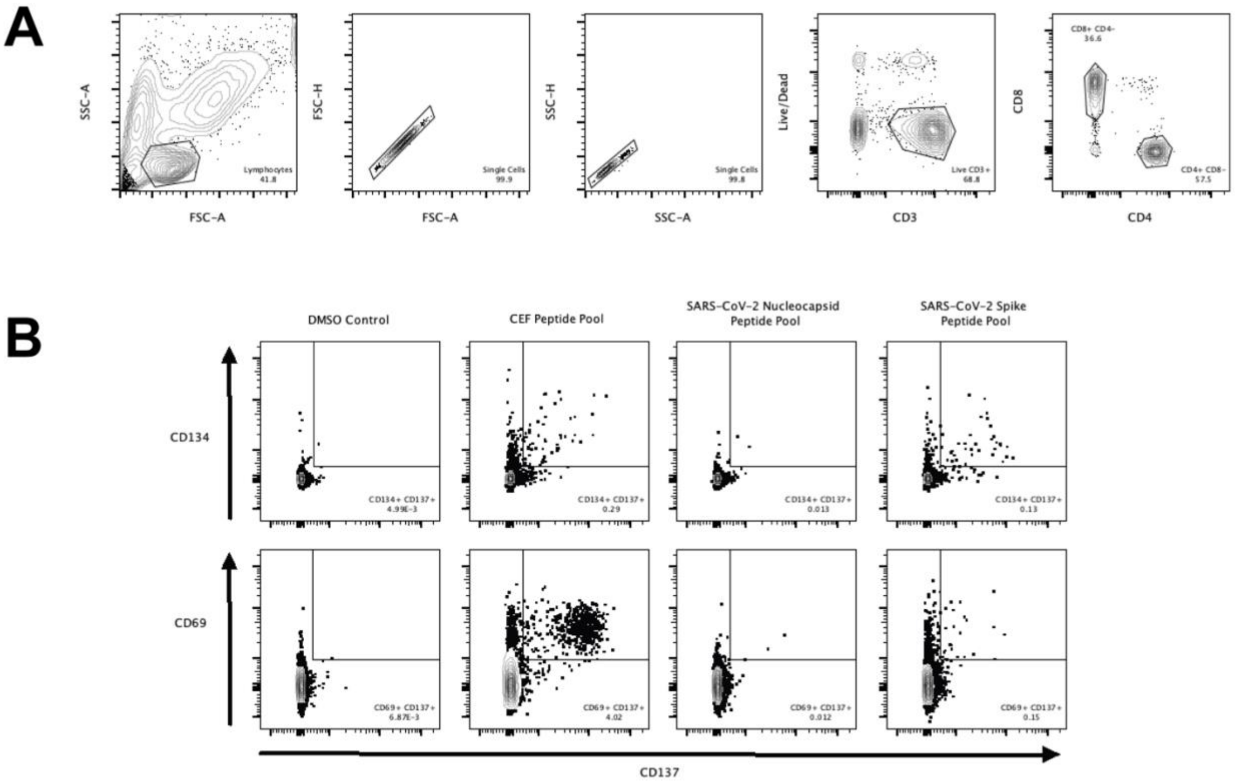

**Figure S2. T cell responses to CMV and CEF peptide pools among those with different SARS-CoV-2 antigen exposure.**

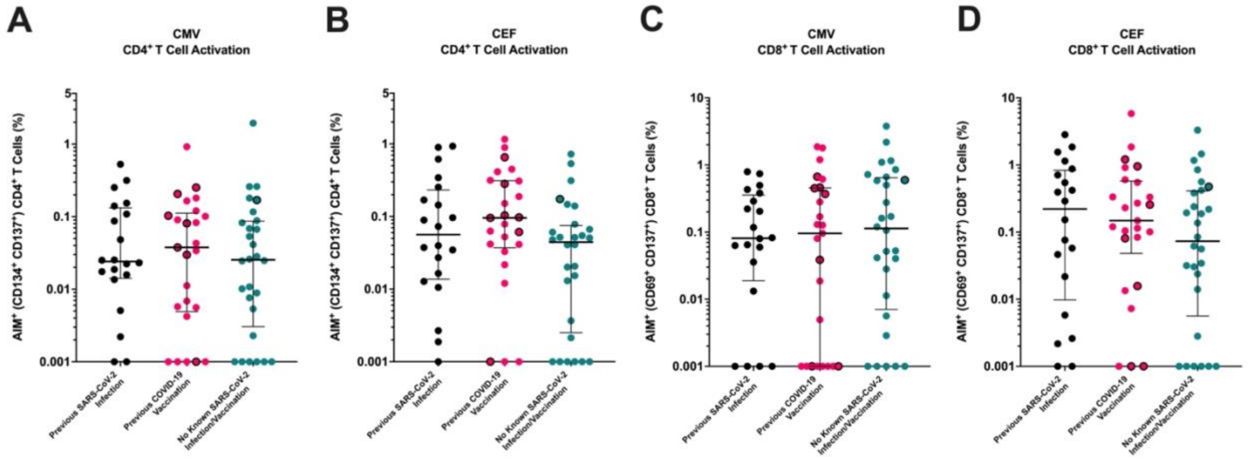

**Figure S3. Alternative analysis of T cell responses to CoV-derived peptide pools among those with different SARS-CoV-2 antigen exposure.**

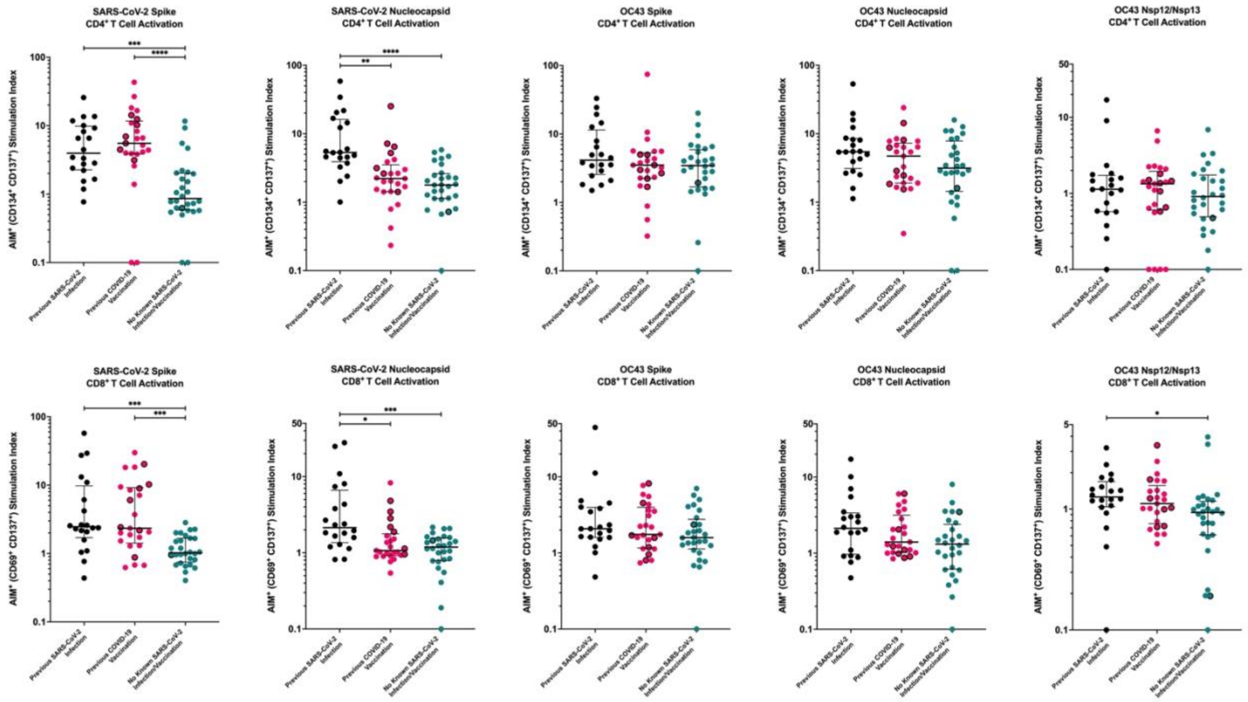

**Table S1. Demographics of the three groups in the retrospective cohort analyses.**

| | Prior SARS-CoV-2 infection (n = 501) | Prior COVID-19 vaccine / No SARS-CoV-2 infection (n = 1,565) | No prior SARS-CoV-2 exposure (n = 2,869) | p-value[A] |
|---|---|---|---|---|
| **Age, median (IQR)** | 53 (36 – 67) | 59 (43 – 71) | 47 (32 – 62) | <0.0001[B] |
| **Male** | 266 (53) | 737 (47) | 1366 (48) | 0.0524 |
| **Race/Ethnicity[C]** | | | | |
| **Black** | 264 (53) | 776 (50) | 1,610 (56) | |
| **White** | 101 (20) | 464 (30) | 629 (22) | <0.0001 |
| **Hispanic/Latino** | 124 (25) | 261 (17) | 501 (17) | |
| **Other/missing** | 20 (4) | 82 (5) | 156 (5) | |
| **Diabetes mellitus** | 143 (29) | 442 (28) | 551 (19) | <0.0001 |
| **Heart disease[D]** | 67 (13) | 217 (14) | 209 (7) | <0.0001 |
| **Lung disease[E]** | 154 (31) | 510 (33) | 717 (25) | <0.0001 |
| **CKD[F]** | 40 (8) | 116 (7) | 126 (4) | <0.0001 |
| **HIV[G]** | 22 (4) | 56 (4) | 74 (3) | 0.0369 |
| **Cancer** | 41 (8) | 171 (11) | 176 (6) | <0.0001 |
| **Smoking** | | | | |
| **Never** | 234 (47) | 648 (41) | 1,332 (46) | |
| **Current** | 107 (21) | 349 (22) | 677 (24) | <0.0001 |
| **Former** | 122 (24) | 425 (27) | 490 (17) | |
| **Missing** | 38 (8) | 143 (9) | 370 (13) | |
| **Number of co-morbidities[H]** | | | | |
| **0** | 206 (41) | 625 (40) | 1,541 (54) | <0.0001 |
| **1** | 179 (36) | 530 (34) | 921 (32) | |
| **≥2** | 116 (23) | 410 (26) | 407 (14) | |
| **Level of care[I]** | | | | |
| **Ambulatory** | 10 (2) | 26 (2) | 42 (1) | 0.6465 |
| **Hospital** | 491 (98) | 1,539 (98) | 2,827 (99) | |

Data shows number and percent unless otherwise indicated.
[A] Chi-square test unless otherwise indicated
[B] Kruskal-Wallis test and Dunn's multiple comparison test.
[C] As specified in the EMR; an individual may be in more than one category
[D] Heart disease includes coronary artery disease and/or congestive heart failure
[E] Lung disease includes chronic obstructive pulmonary disease and/or asthma
[F] Chronic kidney disease
[G] Human immunodeficiency virus
[H] Number of comorbidities accounts for diabetes mellitus, heart disease, lung disease, chronic kidney disease, HIV, and cancer
[I] Level of medical care at the time of the CRP-PCR test. Hospital includes an emergency department visit.

**Table S2. Demographics of the individuals with collected blood specimens.**

| | Prior SARS-CoV-2 infection / No COVID-19 vaccine (n = 20) | Prior COVID-19 vaccine / No SARS-CoV-2 infection (n = 25) | No prior SARS-CoV-2 exposure (n = 28) | p-value[A] |
|---|---|---|---|---|
| **Age, median (IQR)** | 53 (48 – 59) | 62 (59 – 70) | 57 (45 – 65) | 0.0232[B] |
| **Male** | 8 (40) | 17 (68) | 12 (43) | 0.1003 |
| **Race/Ethnicity[C]** | | | | 0.4878 |
| **Black** | 14 (70) | 12 (48) | 16 (57) | |
| **White** | 3 (15) | 10 (40) | 8 (29) | |
| **Hispanic/Latino** | 3 (15) | 3 (12) | 4 (14) | |
| **Other/missing** | 0 (0) | 0 (0) | 0 (0) | |
| **Diabetes mellitus** | 5 (25) | 9 (36) | 10 (36) | 0.6788 |
| **Heart disease[D]** | 3 (15) | 10 (40) | 11 (39) | 0.1359 |
| **Lung disease[E]** | 5 (25) | 7 (28) | 10 (36) | 0.6982 |
| **CKD[F]** | 1 (5) | 2 (8) | 1 (4) | 0.7740 |
| **HIV[G]** | 7 (35) | 4 (16) | 4 (14) | 0.1697 |
| **Cancer** | 1 (5) | 2 (8) | 0 (0) | 0.3328 |
| **Smoking** | | | | 0.0685 |
| **Never** | 11 (55) | 11 (44) | 5 (18) | |
| **Current** | 4 (20) | 5 (20) | 12 (43) | |
| **Former** | 5 (25) | 9 (36) | 11 (39) | |
| **Number of co-morbidities[H]** | | | | 0.6494 |
| **0** | 3 (15) | 5 (20) | 6 (21) | |
| **1** | 12 (60) | 10 (40) | 11 (39) | |
| **≥2** | 5 (25) | 10 (40) | 11 (39) | |
| **Days since last SARS-CoV-2 spike exposure, median (IQR)** | 245 (68-308) | 70 (19-98) | - | 0.0017[I] |

Data shows number and percent unless otherwise indicated.
[A] Chi-square test unless otherwise indicated.
[B] Kruskal-Wallis test.
[C] As specified in the EMR; an individual may be in more than 1 category
[D] Heart disease includes coronary artery disease and/or congestive heart failure
[E] Lung disease includes chronic obstructive pulmonary disease and/or asthma
[F] Chronic kidney disease
[G] Human immunodeficiency virus
[H] Number of comorbidities accounts for diabetes mellitus, heart disease, lung disease, chronic kidney disease, HIV, and cancer
[I] Mann-Whitney U Test

**Table S3: Multi-variable linear regression analysis for predictors of HCoV-OC43 spike binding antibodies.**

| Variable | Estimate (β) | 95% confidence interval | p-value |
|---|---|---|---|
| Intercept | 5.5604 | 5.3999 to 5.7217 | <0.0001 |
| Prior SARS-CoV-2 infection | 0.2586 | 0.0545 to 0.4626 | 0.0140[A] |
| HIV | -0.1613 | -0.3867 to 0.0639 | 0.1580 |
| Hypertension | -0.1343 | -0.3197 to 0.0511 | 0.1530 |

[A] β = 0.2261 and p = 0.0340 after excluding eight with possible occult SARS-CoV-2 infection.

**Table S4. HCoV-OC43 nsp12/nsp12 peptides.**

| Protein (Amino Acid Position) | HCoV-OC43 Peptide | Corresponding SARS-CoV-2 Peptide | % Identity | % Similarity | Suspected HLA Activity |
|---|---|---|---|---|---|
| nsp12 (4393-4407) | SGLSTDVQLRAFDIY | TGTSTDVVYRAFDIY | 73.33 | 80.00 | HLA-B*15:01 |
| nsp12 (4404-4419) | FDIYNASVAGIGLHLK | FDIYNDKVAGFAKFLK | 62.50 | 68.75 | HLA-A*11:01, HLA-A*24:02, HLA-A*32:01 |
| nsp12 (4674-4688) | HCANFNILFSMVLPN | HCANFNVLFSTVFPP | 73.33 | 80.00 | HLA-DQB1*02:02, HLA-DQB1*06:02, HLA-DRB1*07:01 |
| nsp12 (4683-4698) | SMVLPNTCFGPLVRQI | STVFPPTSFGPLVRKI | 68.75 | 68.75 | HLA-A*24:02, HLA-B*57:01 |
| nsp12 (4889-4903) | QDEIYAYTKRNVLPT | QDALFAYTKRNVIPT | 73.33 | 93.33 | HLA-B*51:01, HLA-DRB1*07:01, HLA-DRB1*15:01, HLA-DPA1*02:02/DPB1*05:01 |
| nsp12 (4899-4913) | NVLPTLTQMNLKYAI | NVIPTITQMNLKYAI | 86.67 | 100.00 | HLA-A*01:01, HLA-A*02:06, HLA-B*07:02, HLA-C*07:01 |
| nsp12 (4909-4923) | LKYAISAKNRARTVA | LKYAISAKNRARTVA | 100.00 | 100.00 | HLA-A*68:01, HLA-C*06:02 |
| nsp12 (4944-4958) | IAATRGVPVVIGTTK | IAATRGATVVIGTSK | 80.00 | 93.33 | HLA-A*11:01, HLA-B*51:01, HLA-DQB1*03:03, HLA-DRB1*07:01 |
| nsp12 (4984-4998) | YPKCDRAMPNLLRIV | YPKCDRAMPNMLRIM | 86.67 | 86.67 | **HLA class II** |
| nsp12 (4989-5003) | RAMPNLLRIVSSLVL | RAMPNMLRIMASLVL | 80.00 | 80.00 | HLA-B*07:02, HLA-B*08:01, HLA-C*07:02, HLA-DQB1*06:02, HLA-DRB1*15:01 |
| nsp12 (4997-5011) | IVSSLVLARKHETCC | IMASLVLARKHTTCC | 80.00 | 80.00 | HLA-A*33:01, HLA-A*68:01 |
| nsp12 (5014-5028) | SDRFYRLANECAQVL | SHRFYRLANECAQVL | 93.33 | 93.33 | HLA-DRB1*01:01, HLA-DRB1*08:01, HLA-A*02:01, HLA-A*02:03, HLA-A*02:06 |
| nsp12 (5055-5069) | ANSVFNICQAVSANV | ANSVFNICQAVTANV | 93.33 | 100.00 | HLA-A*02:03, HLA-A*02:06, HLA-A*68:02 |
| nsp12 (5064-5078) | AVSANVCALMSCNGN | AVTANVNALLSTDGN | 66.67 | 80.00 | **HLA-DRB3*02:02, HLA-DRB1*13:02** |

| | | | | | |
|---|---|---|---|---|---|
| nsp12 (5103-5118) | DSTFVTEYYEFLNKHF | DTDFVNEFYAYLRKHF | 56.25 | 75.00 | HLA-A*01:01, HLA-A*02:01, HLA-A*24:02, HLA-A*33:01, HLA-B*08:01 |
| nsp12 (5181-5195) | HTMLVKMDGDDVYLP | HTMLVKQGDDYVYLP | 73.33 | 73.33 | HLA-A*02:01, HLA-B*15:01 |
| nsp12 (5190-5204) | DDVYLPYPNPSRILG | DYVYLPYPDPSRILG | 86.67 | 93.33 | HLA-A*24:02, HLA-B*07:02, HLA-B*51:01 |
| nsp12 (5219-5234) | LLIERFVSLAIDAYPL | LMIERFVSLAIDAYPL | 93.75 | 93.75 | HLA-DQA1*02:01, HLA-DQA1*05:01, HLA-DQA1*01:01, HLA-DQA1*01:03, HLA-DQB1*03:01, HLA-DQB1*06:01, HLA-DQB1*02:02, HLA-DQB1*03:02, HLA-DQB1*04:02, HLA-DQB1*05:01, HLA-DQB1*06:04, HLA-DRB1*01:01, HLA-DRB1*01:03, HLA-DRB1*04:04, HLA-DRB1*07:01, HLA-DRB1*08:03, HLA-DRB1*10:01, HLA-DRB1*12:02, HLA-DRB1*13:02, HLA-DRB1*15:01, HLA-A*02:01, HLA-B*35:01 |
| nsp12 (5229-5243) | IDAYPLVYHENEEYQ | IDAYPLTKHPNQEYA | 66.67 | 73.33 | **HLA class II** |
| nsp12 (5237-5250) | HENEEYQKVFRVYL | HPNQEYADVFHLYL | 57.14 | 78.57 | HLA-A*29:02, HLA-B*44:02, HLA-B*44:03, HLA-C*07:02 |
| nsp12 (5244-5258) | KVFRVYLAYIKKLYN | DVFHLYLQYIRKLHD | 53.33 | 80.00 | HLA-DRB1*04:04, HLA-DRB1*13:02, HLA-C*07:02 |
| nsp13 (5425-5439) | ERLKLFAAETQKATE | ERLKLFAAETLKATE | 93.33 | 93.33 | HLA-A*03:01 |
| nsp13 (5459-5473) | ELILSWEIGKVKPPL | ELHLSWEVGKPRPPL | 73.33 | 86.67 | - |
| nsp13 (5469-5483) | VKPPLNKNYVFTGYH | PRPPLNRNYVFTGYR | 73.33 | 93.33 | - |
| nsp13 (5504-5518) | NGVYYRATTTYKLSV | DAVVYRGTTTYKLNV | 66.67 | 86.67 | HLA-A*11:01, HLA-A*24:02 |

| | | | | | |
|---|---|---|---|---|---|
| nsp13 (5513-5527) | TYKLSVGDVFVLTSH | TYKLNVGDYFVLTSH | 86.67 | 86.67 | HLA-A*02:01 |
| nsp13 (5682-5697) | VINARIRAKHYVYIGD | VVNARLRAKHYVYIGD | 87.50 | 100.00 | HLA-A*11:01 |
| nsp13 (5693-5707) | VYIGDPAQLPAPRVL | VYIGDPAQLPAPRTL | 93.33 | 93.33 | HLA-A*24:02, HLA-C*07:01 |
| nsp13 (5853-5867) | NVNRFNVAITRAKKG | NVNRFNVAITRAKVG | 93.33 | 93.33 | HLA-DRB1*08:03, HLA-DRB1*15:01, HLA-DQA1*05:01, HLA-DQB1*03:01, HLA-DQA1*01:02, HLA-DQB1*06:02 |

**Table S5: Multi-variable linear regression analysis for predictors of HCoV-OC43 nsp12/nsp13 CD8+ T cell responses.**

| Variable | Estimate (β) | 95% confidence interval | p-value |
|---|---|---|---|
| Intercept | -0.0544 | -0.1490 to 0.0402 | 0.2555 |
| Prior SARS-CoV-2 infection | 0.0714 | 0.0223 to 0.1198 | 0.0040 [A] |
| Age > 50 years | 0.00180 | -0.0199 to 0.0742 | 0.2530 |

[A] β = 0.0799 and p = 0.0010 after excluding eight with possible occult SARS-CoV-2 infection
